# The Cul3 ubiquitin ligase engages Insomniac as an adaptor to impact sleep and synaptic homeostasis

**DOI:** 10.1101/689471

**Authors:** Qiuling Li, Kayla Y. Lim, Raad Altawell, Faith Verderose, Xiling Li, Wanying Dong, Joshua Martinez, Dion Dickman, Nicholas Stavropoulos

## Abstract

Mutations of the Cullin-3 (Cul3) E3 ubiquitin ligase are associated with autism and schizophrenia, neurological disorders characterized by sleep disturbances and altered synaptic function. Cul3 engages dozens of adaptor proteins to recruit hundreds of substrates for ubiquitination, but the adaptors that impact sleep and synapses remain ill-defined. Here we implicate Insomniac (Inc), a conserved protein required for normal sleep and synaptic homeostasis in *Drosophila*, as a Cul3 adaptor. Inc binds Cul3 in vivo, and mutations within the N-terminal BTB domain of Inc that weaken Inc-Cul3 associations impair Inc activity, indicating that Inc function requires binding to the Cul3 complex. Deletion of the conserved C-terminus of Inc does not alter Cul3 binding but abolishes Inc activity in the context of sleep and synaptic homeostasis, suggesting that the C-terminal domain of Inc is a substrate recruitment domain. Mutation of a conserved, disease-associated arginine in the Inc C-terminus also abolishes Inc function, suggesting that this residue is vital for recruiting Inc targets. Inc levels are negatively regulated by Cul3 in neurons, consistent with Inc degradation by autocatalytic ubiquitination, a hallmark of Cullin adaptors. These findings link Inc and Cul3 in vivo and indicate that Cul3-Inc complexes are essential for normal sleep and synaptic function. Furthermore, these results indicate that dysregulation of conserved substrates of Cul3-Inc complexes may contribute to altered sleep and synaptic function in autism and schizophrenia associated with *Cul3* mutations.

**Author Summary:** *Cul3* is a highly conserved gene important for brain development and function. *Cul3* mutations are a risk factor for autism and schizophrenia, neurological disorders associated with disturbed sleep and changes in neuronal synapses. A key challenge in understanding how *Cul3* impacts brain function is elucidating the downstream molecular pathways. Cul3 is a ubiquitin ligase that assembles with dozens of adaptor proteins, which in turn recruit specific protein substrates for ubiquitination. Identifying and characterizing Cul3 adaptors in the nervous system is thus a critical step in understanding Cul3 function. Because Cul3 and its adaptors are conserved through evolution, simpler organisms including the fruit fly *Drosophila* provide powerful systems for identifying and characterizing Cul3 adaptors. We found that the Insomniac (Inc) protein has the properties of a Cul3 adaptor that impacts sleep and synaptic function in *Drosophila*. These results suggest that human proteins related to Inc may be relevant for changes in sleep and synaptic function in autism and schizophrenia associated with reduced Cul3 activity.

## Introduction

The Cullin-3 (Cul3) E3 ubiquitin ligase is broadly expressed in the brain and is an important regulator of nervous system development and function. In humans, *Cul3* mutations are linked to autism, its associated sleep disturbances, and to schizophrenia [1–7]. Mice bearing *Cul3* mutations exhibit alterations in neurogenesis, dendritic development, synaptic transmission, and glutamate receptor levels, as well as social deficits relevant to autism spectrum disorder [8–14]. *Cul3* is highly conserved, and reduced activity of *Cul3* in worms and flies causes similar phenotypes within the nervous system, including perturbations in neuronal morphogenesis, glutamate receptor abundance, synaptic homeostasis, and sleep [15–20].

A key challenge in understanding how *Cul3* impacts brain function is elucidating the downstream molecular pathways. Cul3 assembles in a modular fashion with dozens of BTB-domain adaptor proteins that recruit substrates to Cul3 complexes for ubiquitination [21–25]. Which among the known and putative adaptors of Cul3 contribute to its phenotypes—including sleep disturbances and altered synaptic function—remains largely unknown. Though several Cul3 adaptors have been characterized in neurons [16,26–32], many known and putative Cul3 adaptors have yet to be assessed.

*insomniac* (*inc*) encodes a conserved BTB-domain protein that is hypothesized to serve as a Cul3 adaptor [17,18,33]. *inc* null mutations reduce sleep duration and consolidation in *Drosophila* and impair synaptic homeostasis, phenotypes that are recapitulated by reduced *Cul3* activity [17,18,20]. Inc can bind Cul3 when overexpressed in cultured cells [17,18,33], but whether these interactions occur in vivo or are required for the activity of Inc is unknown. Notably, some BTB-domain proteins associate with Cul3 but are unlikely to serve as adaptors [34]. Functional analysis has been essential to distinguish Cul3 adaptors within the BTB superfamily, which also includes GABA_B_ receptor subunits, potassium channels, and transcriptional regulators [35,36]. In the absence of evidence linking Inc and Cul3 in vivo, the role of Inc as a Cul3 adaptor has remained speculative.

Here we implicate Inc as a Cul3 adaptor, using biochemical, behavioral, and electrophysiological studies of Inc mutants. Critically, Inc binds Cul3 in vivo and Inc activity requires the assembly of Inc-Cul3 complexes. Inc function also requires its C-terminus, a conserved domain dispensable for Cul3 associations but vital for Inc activity, consistent with the properties of a substrate recruitment domain. Mutation of a conserved C-terminal arginine abolishes Inc function, suggesting that this residue is vital for recruiting Inc targets. Mutation of the same residue in the human Inc ortholog KCTD17 is associated with myoclonic dystonia [37], consistent with a critical and evolutionarily conserved function for this residue in Inc orthologs. Our results reveal that Cul3-Inc complexes impact sleep and synaptic function, and provide tools to identify targets of Cul3, Inc, and Inc orthologs that impact normal brain function and neurological disorders.

## Results

### The Inc BTB domain mediates Cul3 binding and Inc multimerization

Cul3 adaptors have a modular functional organization, with BTB domains that self-associate and bind Cul3, and distal domains that recruit substrates [21–24]. Inc has two conserved domains, an N-terminal BTB domain and a C-terminal domain unique to Inc and its orthologs [17,38,39] (Fig 1A and S1A Fig). To determine whether Inc has the properties of a Cul3 adaptor, we generated progressive N- and C-terminal deletions of Inc (Fig 1A) and assessed whether these derivatives could bind Inc and Cul3, by expressing epitope-tagged forms of these proteins in *Drosophila* S2 cells and performing co-immunoprecipitations.

**Fig 1.**
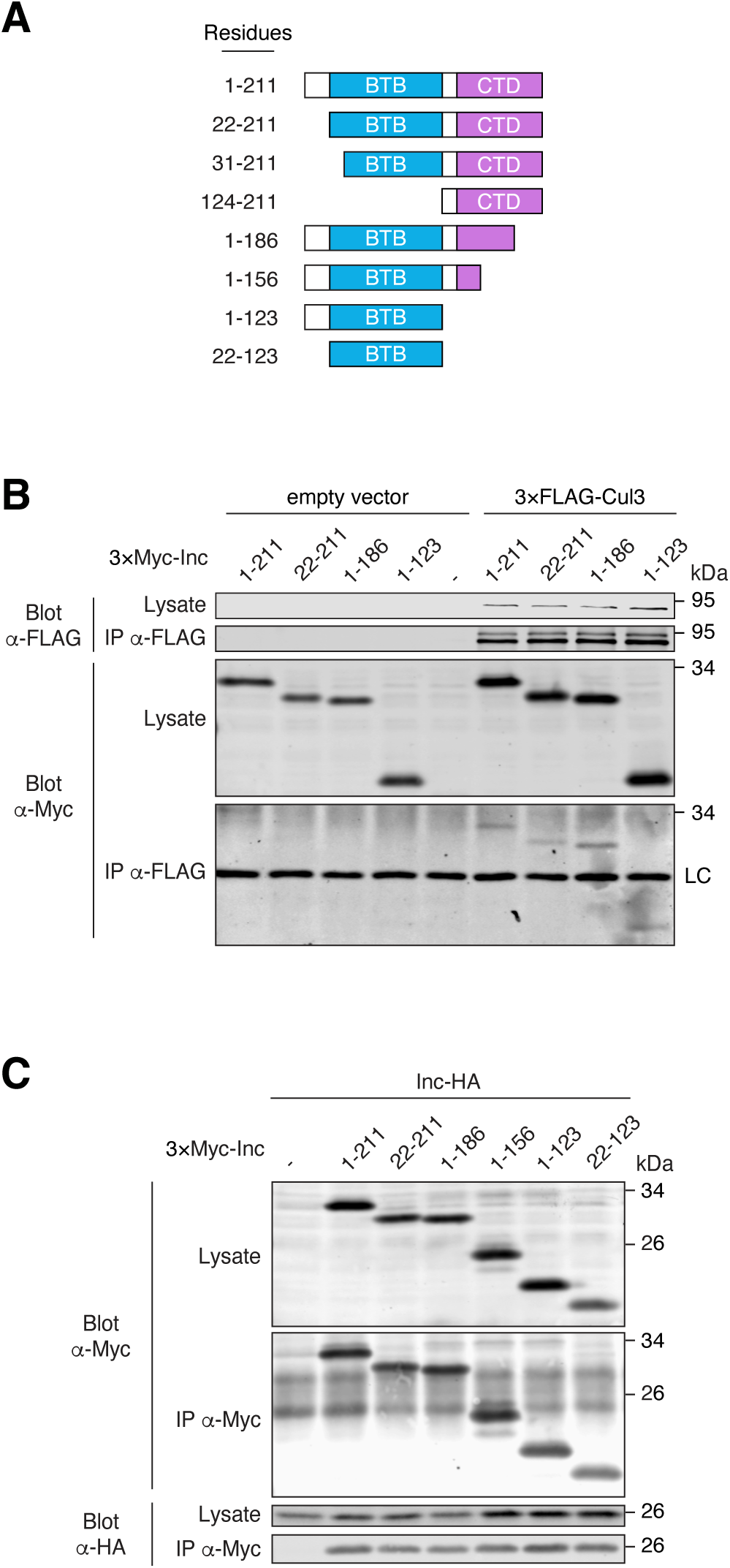
The Inc BTB domain mediates Inc-Cul3 and Inc-Inc interactions. **(A)** Schematic of N- and C-terminally truncated Inc proteins. Residues comprising the conserved BTB domain (22 to 123) and C-terminal domain (CTD) (131 to 211) are indicated. **(B and C)** Co-immunoprecipitation of 3×Myc-tagged Inc or Inc truncation mutants with 3×FLAG-Cul3 **(B)** or Inc-HA **(C)** from transiently transfected S2 cells. LC, immunoglobulin light chain.

Deleting Inc residues preceding the BTB domain (Inc^22-211^) did not alter Inc stability or Inc-Cul3 interactions (Fig 1B and S2 Fig), indicating that the N-terminus of Inc is dispensable for Cul3 binding. In contrast, removing part of the BTB domain (Inc^31-211^) or the entire BTB domain (Inc^124-211^) significantly destabilized Inc (S2 Fig), and low steady-state levels of these proteins precluded conclusions about their ability to bind Cul3. Inc proteins lacking 25 C-terminal residues (Inc^1-186^) or all residues following the BTB domain (Inc^1-123^) were stable and associated with Cul3 similarly to full-length Inc. An Inc mutant lacking most of the Inc C-terminal domain (Inc^1-156^) was stable and interacted with Cul3 more strongly than wild-type Inc and other Inc deletions (S3A Fig). Thus, the Inc C-terminus is dispensable for interaction with Cul3. The Inc BTB domain alone (Inc^22-123^) was sufficient to bind Cul3, albeit more weakly than wild-type Inc (S3B Fig); because deleting the same N-terminal residues from full length Inc slightly weakened Cul3 binding (Fig 1B; Inc^22-211^ versus Inc^1-211^), these results suggest that Inc N-terminal residues stabilize Cul3 interactions mediated chiefly by the Inc BTB domain.

Cul3 adaptors homomultimerize through their BTB domains, and adaptor-mediated oligomerization of Cul3 complexes can enhance substrate ubiquitination [40]. We therefore assessed whether Inc self-association is mediated by its BTB domain. N-terminal truncations that preserve the Inc BTB domain (Inc^22-211^) did not alter Inc-Inc associations (Fig 1C). Similarly, Inc proteins truncated C-terminally (Inc^1-186^, Inc^1-156^, and Inc^1-123^) associated with Inc normally, indicating that the Inc C-terminus is dispensable for Inc multimerization (Fig 1C). The Inc BTB domain (Inc^22-123^) alone was sufficient to bind Inc (Fig 1C). Thus, the Inc BTB domain is sufficient to mediate both Inc multimerization and Cul3 binding, as expected of a Cul3 adaptor.

### Inc point mutants define a surface of the Inc BTB domain that binds Cul3

To determine whether Inc function in vivo requires binding to Cul3 and Inc homomultimerization, we sought to identify Inc point mutants that selectively perturb Inc-Cul3 and Inc-Inc interactions. Our mutagenesis of Inc was informed by the structures of homodimeric BTB-MATH and BTB-BACK-Kelch adaptors bound to Cul3 [40–42], and by the structure of KCTD5, a human Inc ortholog that forms homopentamers [39]. Homology modeling and light scattering measurements suggest that Inc forms a homopentamer similar to that of KCTD5 [43,44]. We selected eight conserved residues in the Inc BTB domain whose equivalents in KCTD5 reside on a surface suggested to bind Cul3 [43,45] (S1A Fig). We mutated these residues to alanine (F47A, D57A, D61A, F105A, N107A), to oppositely charged residues (R50E, E104K), or altered their hydrophobicity (Y106F), and assessed the effects on Inc-Cul3 and Inc-Inc binding. Most of these mutants did not alter Inc-Cul3 or Inc-Inc interactions, while one mutant (R50E) impaired both associations (S1 Table). Two mutants, F47A and F105A (Fig 2A), weakened interactions with Cul3 (Fig 2B) but did not significantly affect Inc self-association or stability (Fig 2C and S4 Fig). Similarly, an F47A/F105A double mutant was stable and selectively impaired Cul3 binding (Fig 2B and 2C and S4 Fig). The analogous phenylalanines in KCTD5 cluster near the interface of adjacent subunits, defining a surface for Cul3 binding (S1B Fig).

**Fig 2.**
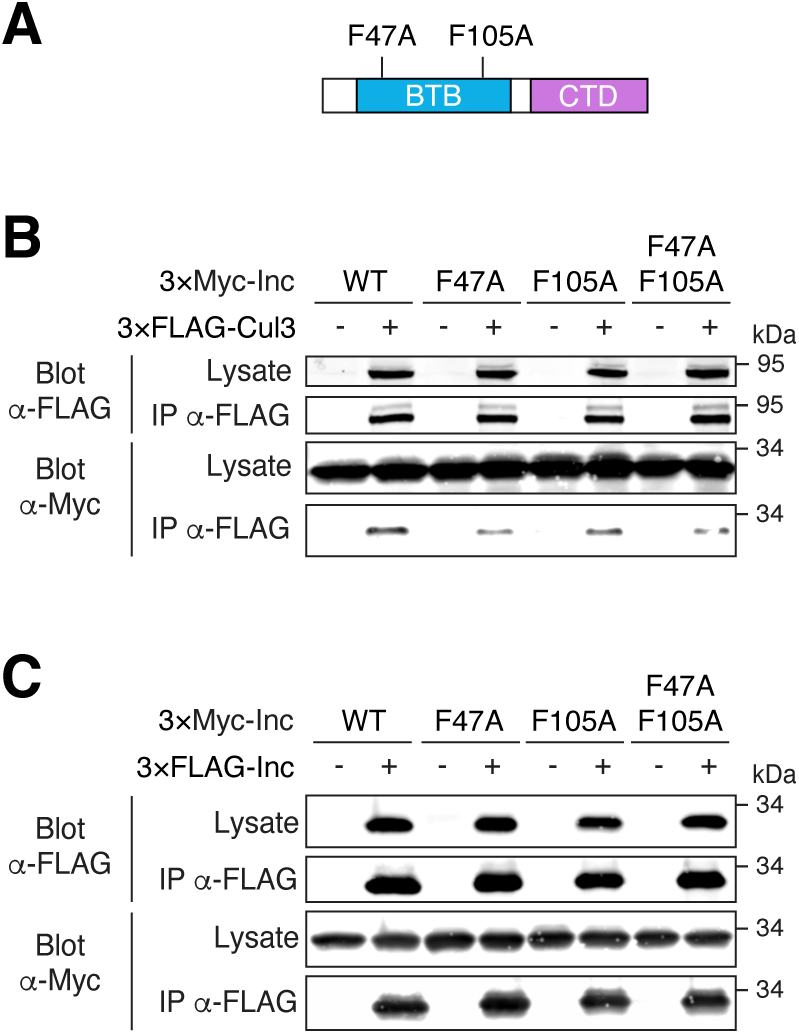
Identification of Inc point mutants that impair Inc-Cul3 interactions. **(A)** Schematic of Inc point mutants within the BTB domain. **(B-C)** Co-immunoprecipitation of 3×Myc-tagged Inc or Inc point mutants with 3×FLAG-Cul3 **(B)** or 3×FLAG-Inc **(C)** from transiently transfected S2 cells.

To identify Inc mutants that impair its multimerization, we selected seven conserved residues whose equivalents in KCTD5 engage in polar and charged interactions between adjacent subunits [39] (S1A and S1C Fig). Mutating these residues individually or in double mutant combinations to alanine (T36A, D71A, D73A, N82A) or to oppositely charged residues (R85E, K88D, E101K) did not significantly alter Inc self-association (S2 Table). A triple mutant (T36A/D71A/R85E) predicted to disrupt interactions of Inc subunits with both flanking neighbors in a putative Inc homopentamer exhibited significantly reduced stability (S4 Fig and S2 Table), precluding conclusions about its ability to multimerize and bind Cul3. Because this triple mutant was also unstable when expressed in vivo, we focused further analysis on Inc mutants that selectively perturb Cul3 interactions.

### Inc function in vivo requires the assembly of Inc-Cul3 complexes

To determine whether Inc impacts sleep as a Cul3 adaptor, we assessed whether Inc mutants that weaken Cul3 binding impair Inc activity in vivo. We generated UAS transgenes expressing 3×FLAG-tagged Inc, Inc^F47A^, Inc^F105A^, and Inc^F47A/F105A^, integrating them at the same genomic site to compare their activity. When expressed with *inc-Gal4*, a driver that rescues *inc* mutants when used to restore *inc* expression [17,33], Inc and Inc mutants were similarly stable in both *inc^1^* and *inc^+^* animals (Fig 3A and S5 Fig). Because Inc mutants deficient for Cul3 binding might sequester substrates and function as dominant negatives, we tested whether these mutants could alter sleep when expressed in neurons. Neuronal expression of Inc^F47A^, Inc^F105A^, or Inc^F47A/F105A^ did not alter sleep (S6A Fig), in contrast to neuronal *inc* RNAi [17,18]. Similarly, expression of these mutants using *inc-Gal4* did not perturb sleep (S6D Fig). Thus, these Inc mutants do not antagonize endogenous Inc, possibly because they retain some ability to bind Cul3 (Fig 2B). Finally, we tested whether Inc function in vivo requires the assembly of Inc-Cul3 complexes. If so, Inc mutants that weaken Cul3 binding should impair *inc* rescue. Whereas wild-type Inc expressed with *inc-Gal4* fully rescued daily sleep duration in *inc^1^*mutants, Inc^F47A^ provided only a partial rescue, and Inc^F105A^ and Inc^F47A/F105A^ failed to restore a significant amount of sleep (Fig 3B and 3C), indicating that Inc activity requires binding to the Cul3 complex. For other sleep parameters, rescue was more strongly impaired for Inc^F105A^ and Inc^F47A/F105A^ than for Inc^F47A^ (S7A–D Fig), suggesting that F105 is particularly critical for Cul3 binding in vivo. All rescuing activity of UAS transgenes was Gal4-dependent, as indicated by the short sleep of *inc^1^* mutants bearing only UAS transgenes (S8 Fig). Collectively, these results indicate that Inc function in vivo requires the assembly of Inc-Cul3 complexes and that their disruption inhibits sleep.

**Fig 3.**
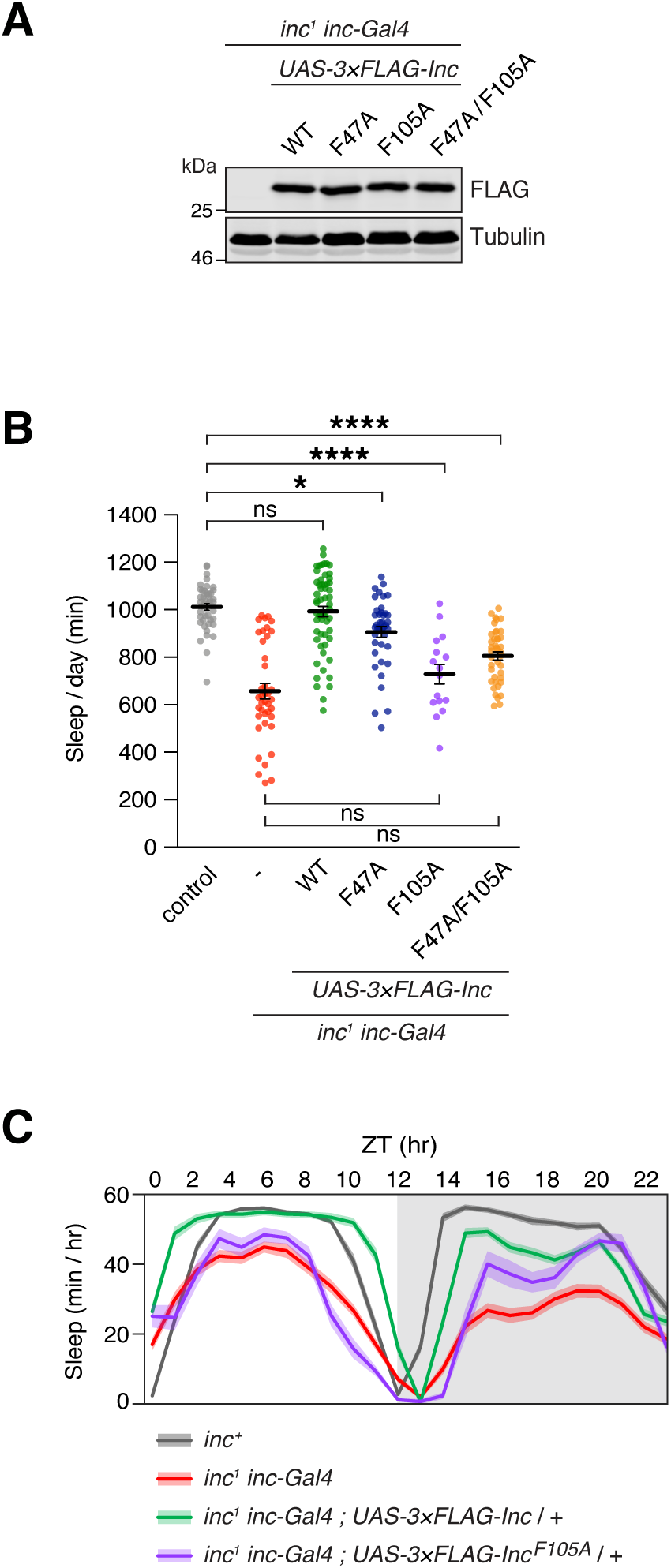
Inc-Cul3 binding is required for Inc activity in vivo. In vivo analysis of *inc^1^ inc-Gal4* animals expressing 3×FLAG-tagged Inc or Inc point mutants. **(A)** Immunoblot of male whole animal lysates. **(B)** Total sleep duration. Mean ± SEM is shown. N = 16-58; Kruskal-Wallis, p<0.0001 and Dunn’s tests; *p < 0.05; ****p < 0.0001; ns, not significant (p > 0.5). **(C)** Average daily sleep traces of indicated genotypes from **(B)**. Shading represents SEM. For all panels, animals are heterozygous for UAS transgenes.

### The Inc C-terminus is vital for Inc activity in vivo and has the properties of a substrate recruitment domain

Although the Inc C-terminus is dispensable for Inc-Inc and Inc-Cul3 associations (Fig 1B and 1C), its evolutionary conservation suggests a critical function. The modular structure of Cul3 adaptors suggests that if Inc serves as a Cul3 adaptor, its C-terminus would bind and recruit substrates to Inc-Cul3 complexes. We therefore assessed the activity of C-terminally truncated Inc mutants in vivo. 3×FLAG-tagged Inc^1-186^, Inc^1-156^, and Inc^1-123^ were expressed at similar levels under *inc-Gal4* control (Fig 4A), indicating that they are stable in vivo, as in cultured cells (Fig 1B and 1C). Expression of C-terminally truncated Inc mutants neuronally with *elav-Gal4* or with *inc-Gal4* did not alter sleep, indicating that they do not inhibit endogenous *inc* function (S6B Fig and S6D Fig). We next tested whether C-terminally truncated Inc proteins could restore sleep to *inc* mutants. Wild-type Inc expressed with *inc*-*Gal4* fully rescued sleep in *inc^1^* mutants (Fig 4B and 4C). In contrast, C-terminal truncations of Inc curtailed or abolished Inc function. Inc^1-186^ partially rescued *inc* mutants, while animals expressing Inc^1-156^ or Inc^1-123^ slept indistinguishably from *inc* mutants, indicating that Inc^1-156^ and Inc^1-123^ lacked activity in vivo (Fig 4B and 4C and S9A–D Fig). Thus, the Inc C-terminus is vital for Inc function. The ability of C-terminally truncated Inc proteins to multimerize and bind Cul3 normally (Fig 1B and 1C) is consistent with a modular function for the Inc C-terminus in recruiting targets that impact sleep.

**Fig 4.**
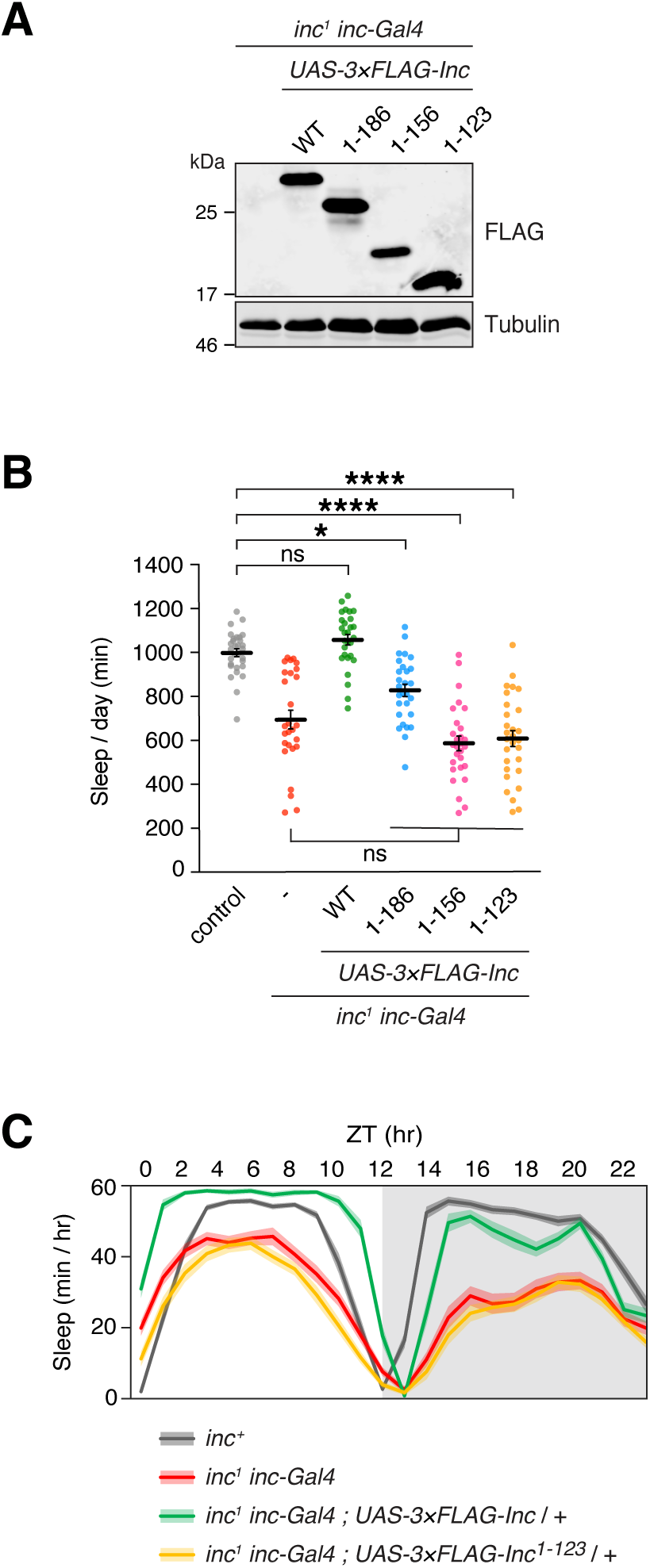
The conserved Inc C-terminus is required for Inc activity in vivo. In vivo analysis of *inc^1^ inc-Gal4* animals expressing 3×FLAG-tagged Inc or Inc truncation mutants. **(A)** Immunoblot of male whole animal lysates. **(B)** Total sleep duration. Mean ± SEM is shown. n = 27-30; Kruskal-Wallis, p < 0.0001 and Dunn’s tests; *p < 0.05; ****p < 0.0001; ns, not significant (p > 0.5). **(C)** Average daily sleep traces of indicated genotypes from **(B)**. Shading represents SEM. For all panels, animals are heterozygous for UAS transgenes.

### Inc-Cul3 binding and the Inc C-terminus are vital for Inc activity at synapses

We next assessed whether Inc-Cul3 associations and the Inc C-terminus are required for Inc function at synapses. At the larval neuromuscular junction, Inc is required for the homeostatic upregulation of presynaptic neurotransmitter release caused by inhibition of postsynaptic glutamate receptors [20]. Application of philanthotoxin, a glutamate receptor antagonist, reduces the amplitude of spontaneous neurotransmission, while homeostatic upregulation of evoked neurotransmission maintains synaptic strength in wild-type animals (Fig 5A and 5B) [46]. Synaptic homeostasis is blocked in *inc* mutants and is restored by Inc expressed post-synaptically in muscle with *BG57-Gal4* [20] (Fig 5A and 5B). We assessed whether Inc^F105A^, which impairs Cul3 associations and rescue of sleep (Figs 2B, 3B, and 3C), could restore synaptic homeostasis to *inc* mutants. In contrast to wild-type Inc, post-synaptic expression of Inc^F105A^ failed to restore synaptic homeostasis to Inc mutants, indicating that Inc-Cul3 associations are required for Inc activity at synapses (Fig 5B). To determine whether the activity of Cul3-Inc complexes requires the Inc C-terminus, we similarly assessed the rescuing activity of Inc^1-123^, which lacks this conserved domain. Inc^1-123^ failed to restore synaptic homeostasis to *inc* mutants (Fig 5A and 5B), indicating that the C-terminus of Inc is essential for Inc activity at synapses. The requirement for Inc-Cul3 associations and the Inc C-terminus for both normal sleep and synaptic plasticity supports the notion that Inc serves as a Cul3 adaptor in vivo.

**Fig 5.**
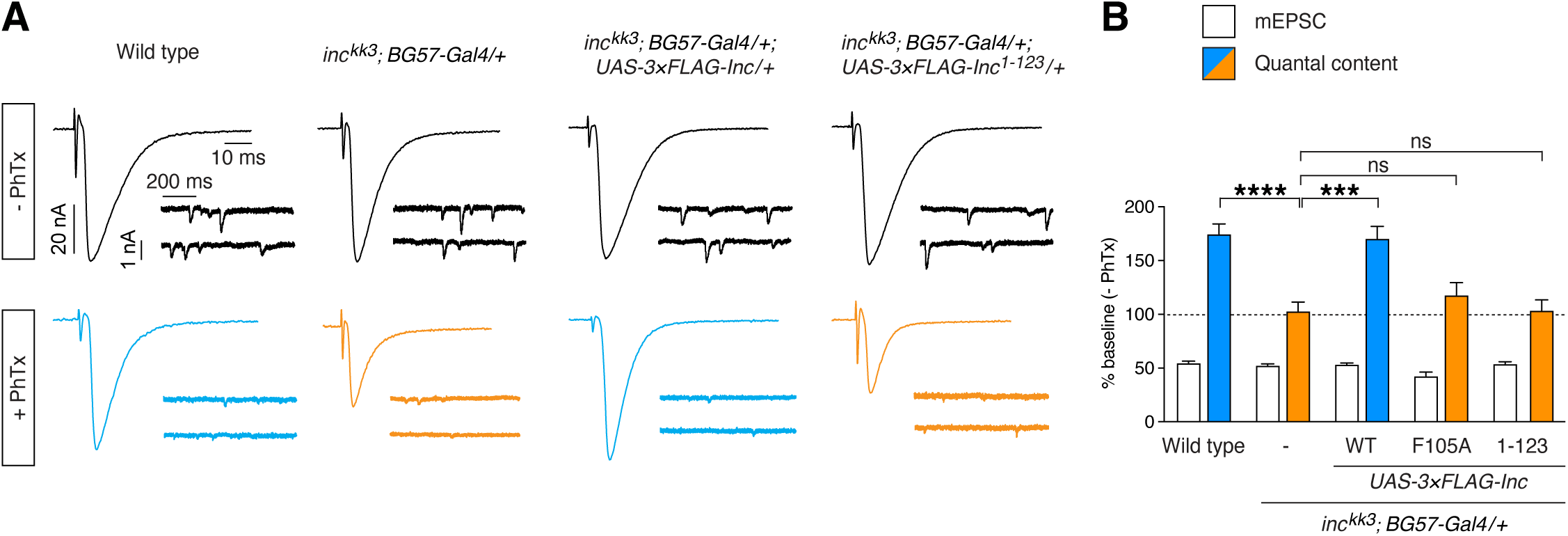
Inc-Cul3 associations and the Inc C-terminus are essential for synaptic homeostasis. **(A)** Spontaneous and evoked electrophysiological traces at the larval NMJ in the presence and absence of philanthotoxin. **(B)** Summary data for electrophysiological recordings in the presence of philanthotoxin normalized to baseline values (absence of PhTx application). Homeostatic plasticity is not expressed in animals bearing Inc^F105A^ or Inc^1-123^. Mean ± SEM is shown. n = 8-27; Kruskal-Wallis, p<0.0001 and Dunn’s tests; ***p < 0.001; ****p < 0.0001; ns, not significant (p > 0.5).

### Mutation of a conserved disease-associated arginine abolishes Inc function and defines a surface likely to recruit substrates

To further dissect Inc function, we mutated conserved C-terminal residues we hypothesized might bind Inc targets (Fig 6A). In particular, a missense mutation of a conserved arginine (R145H) in the human Inc ortholog KCTD17 is associated with myoclonic dystonia, a movement disorder [37]. While the mechanism of KCTD17^R145H^ pathogenesis is unknown, the analogous KCTD5 residue resides on the surface of the C-terminal domain (S1D Fig). We reasoned that this residue might have a conserved function in binding substrates, and we therefore mutated the corresponding Inc residue (Inc^R135H^) to assess its function. We also mutated three Inc residues (N167, Y168, G169) to alanine (Inc^AAA^); the corresponding KCTD5 residues reside in an exposed loop that might recruit substrates (S1D Fig).

**Fig 6.**
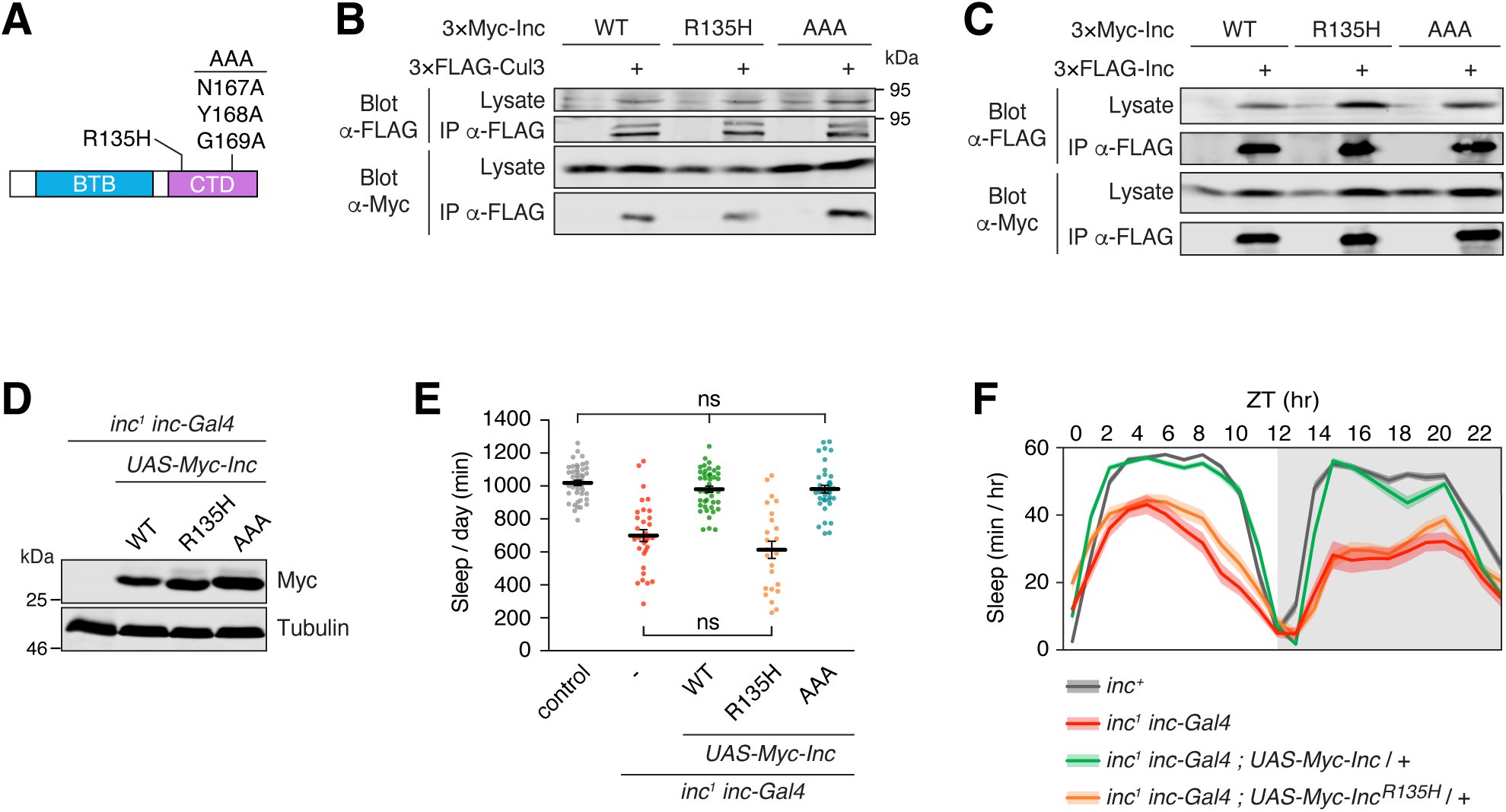
A conserved Inc C-terminal arginine is vital for Inc activity. **(A)** Schematic of Inc C-terminal point mutants. **(B and C)** Co-immunoprecipitation of 3×Myc-tagged Inc or Inc point mutants with 3×FLAG-Cul3 **(B)** or 3×FLAG-Inc **(C)** from transiently transfected S2 cells. **(D)** Immunoblot of lysates prepared from whole male animals expressing 3×Myc-tagged Inc proteins. **(E)** Total sleep duration. Mean ± SEM is shown. n = 24-45; Kruskal-Wallis, p < 0.0001 and Dunn’s tests; ns, not significant (p > 0.05). **(F)** Average daily sleep traces of indicated genotypes from **(E)**. Shading represents SEM. For panels **(D-F)**, animals are heterozygous for UAS transgenes.

In S2 cells, Inc^R135H^ and Inc^AAA^ were stable and associated with Cul3 and Inc indistinguishably from wild-type Inc (Figs 6B and 6C), indicating that these mutations do not alter Inc multimerization or Cul3 binding. In vivo, Inc^R135H^ and Inc^AAA^ were stably expressed with *inc-Gal4* (Fig 6D) and lacked dominant negative activity when expressed in neurons or with *inc-Gal4* (S6C–D Fig). Next, we tested whether Inc^R135H^ and Inc^AAA^ could rescue the sleep deficits of *inc* mutants. When expressed with *inc-Gal4*, Inc^AAA^ rescued *inc^1^* mutants indistinguishably from wild-type Inc, indicating that Inc^AAA^ retains normal activity (Fig 6E and 6F and S8 Fig and S10A–D Fig). In stark contrast, Inc^R135H^ was unable to restore sleep to *inc^1^* mutants, indicating that the R135H mutation abolishes Inc function (Fig 6E and 6F and S10A–D Fig). The complete loss of Inc activity caused by the R135H mutation is indistinguishable from the consequences of deleting the Inc C-terminus (Fig 4B and 4C), suggesting that R135 is essential for the function of the C-terminus and binding Inc targets. These findings furthermore suggest that the analogous KCTD17^R145H^ mutation causes myoclonic dystonia by impairing the recruitment and ubiquitination of KCTD17 targets.

### Inc exhibits properties of a Cul3 adaptor in neurons and is negatively regulated by Cul3

Normal sleep requires the activity of both *inc* and *Cul3* in neurons [17,18]. We performed several experiments to determine whether Inc has the properties of a Cul3 adaptor in vivo. First, we tested whether Inc-Cul3 complexes assemble in vivo, by co-expressing epitope-tagged Inc and Cul3 with *inc-Gal4* and performing co-immunoprecipitations. We observed that Inc and Cul3 associate in vivo (Fig 7A). Second, we tested whether Inc multimerizes in vivo, by assessing the physical interactions of Inc-HA and Myc-Inc co-expressed with *inc-Gal4*. These proteins associated robustly (Fig 7B), indicating that Inc forms homomultimers in vivo. Third, we tested whether Inc is regulated by Cul3. Cul3 adaptors are often regulated by autocatalytic ubiquitination and degradation in Cul3 complexes [22,47–49]. Neuronal RNAi against *Cul3* increased Inc protein levels in head lysates (Fig 7C), indicating that Cul3 negatively regulates Inc abundance. qRT-PCR confirmed that neuronal Cul3 RNAi reduced Cul3 mRNA levels in heads (Fig 7D), as reported previously [50], but did not significantly alter *inc* transcript levels (Fig 7D). These findings indicate that Cul3 negatively regulates Inc by a post-transcriptional mechanism. We conclude that Inc is likely regulated by Cul3-dependent autocatalytic ubiquitination and degradation in neurons, cells through which *inc* influences sleep [17,18]. Together, these results indicate that Inc has the attributes of a Cul3 adaptor in vivo.

**Fig 7.**
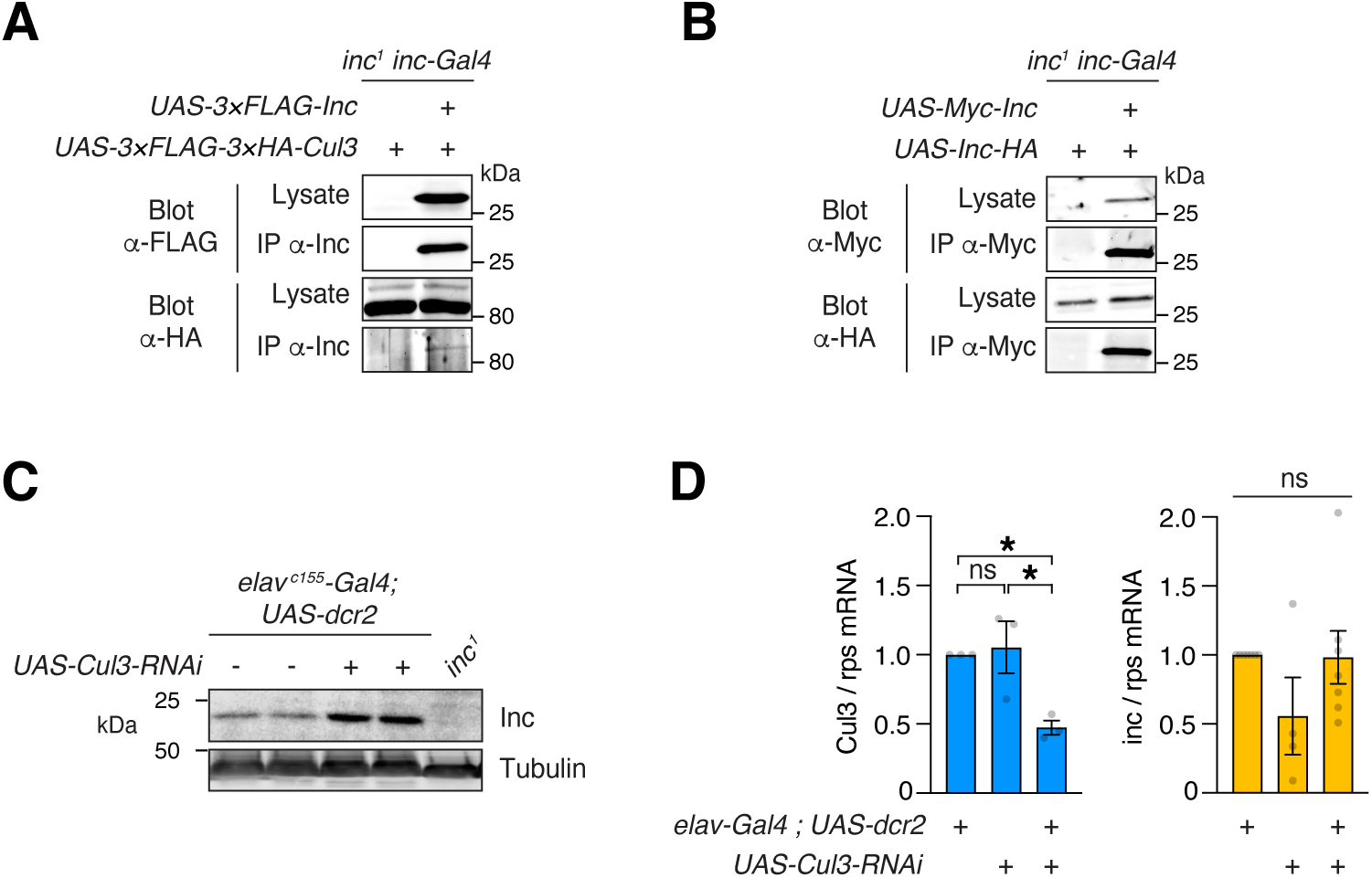
Inc exhibits key properties of a Cul3 adaptor in neurons in vivo. **(A)** Co-immunoprecipitation of 3×FLAG-Inc and 3×FLAG-3×HA-Cul3 from male whole animal lysates. Note immunoprecipitation (IP) was performed with anti-Inc antibody. **(B)** Co-immunoprecipitation of Myc-Inc and Inc-HA from male whole animal lysates. **(C)** Immunoblots of male head lysates. **(D)** qRT-PCR of *Cul3* and *inc* mRNA isolated from male heads. Neuronal *Cul3* RNAi decreases *Cul3* transcript levels but does not significantly alter those of *inc*. Each point represents an independent biological sample measured in technical triplicate. n=3-7. Mean ± SEM is shown. For *Cul3*, one-way ANOVA, p < 0.05 and Tukey tests; for *inc*, Kruskal-Wallis; *p < 0.05; ns, not significant (p > 0.25).

## Discussion

*Cul3* mutations disrupt brain development and physiology and contribute to neurological disorders, including autism and schizophrenia. Understanding the diverse functions of *Cul3* in the nervous system requires characterization of its relevant adaptors and substrates. Our findings implicate Inc as a Cul3 adaptor critical for sleep and synaptic function. We have shown that Inc-Cul3 complexes assemble in vivo, that weakening Inc-Cul3 associations impairs Inc activity, and that Inc has modular functional domains typical of Cul3 adaptors. The conserved C-terminal domain of Inc is vital for Inc activity but dispensable for Inc multimerization and Cul3 binding, as expected of a substrate recruitment domain. Lastly, the negative regulation of Inc abundance by Cul3 in vivo is characteristic of autocatalytic adaptor ubiquitination within the Cul3 complex and subsequent proteolytic degradation.

In addition to demonstrating that Inc is regulated by autocatalytic ubiquitination, the present findings suggest that Inc activity is controlled by additional mechanisms. Inc mutants deficient for Cul3 binding would be expected to sequester substrates but do not perturb sleep when overexpressed. Similarly, C-terminally truncated Inc mutants that cannot recruit substrates might be expected to compete for Cul3 binding and inhibit ubiquitination of Inc substrates, yet these mutants lack dominant negative activity. Consistent with these findings, overexpression of wild-type Inc rescues *inc* mutants in a permissive manner but does not cause gain-of-function phenotypes, for both sleep and synaptic homeostasis [17,20,33]. Inc may thus be regulated by mechanisms that control its ability to bind Cul3, recruit substrates, or catalyze substrate ubiquitination.

Inc and its orthologs are structurally distinct from better characterized Cul3 adaptors of the BTB-MATH and BTB-BACK-Kelch families. In contrast to BTB-MATH and BTB-BACK-Kelch adaptors that form homodimers and bind Cul3 independently [40–42], Inc orthologs form homopentamers that bind Cul3 in radial assemblies [39,43,45,51–53]. Our findings indicate that Cul3 binds to a cleft between adjacent Inc subunits, consistent with a similar homopentameric structure for Cul3-Inc complexes. Because formation of higher order Cul3 complexes can stimulate substrate ubiquitination [40], the pentameric structure of Inc and its orthologs may enable efficient ubiquitination of substrates, including those that associate weakly or transiently with the Cul3 complex.

The analysis of C-terminal Inc truncations and point mutations suggests a mechanism by which Inc and its orthologs bind substrates. Deleting the 25 C-terminal residues of Inc reduces but does not abolish Inc activity, indicating that these residues contribute to substrate binding. The analogous KCTD5 residues are not resolved in crystal structures of KCTD alone [39] (Fig. S1D), suggesting that they are flexible and may become structured upon target binding. Second, mutation of a surface-exposed arginine (R135) abolishes Inc function and is functionally equivalent to deleting the C-terminus. Together, these data support a model in which R135 and the surrounding lateral surface of Inc bind substrates in conjunction with the C-terminal tail of Inc. A recent structure of KCTD5 bound to a substrate provides direct support for this mechanism [53]. Furthermore, this mechanism may be conserved in other Inc orthologs and disrupted by KCT17^R145H^ mutations linked to myoclonic dystonia [37]. While the simplest hypothesis is that Inc binds substrates directly, Inc might also engage co-adaptor proteins, analogous to the mechanism described for the BTB-Kelch adaptor KLHL12 [54].

Our findings do not identify Inc targets that mediate its impact on sleep and synaptic homeostasis in *Drosophila*, nor do they discern whether Inc orthologs contribute to changes in sleep or other *Cul3* phenotypes in mammals. Another important unresolved question is whether heterozygous loss-of-function *Cul3* mutations alter sleep in rodent models, analogous to sleep disturbances reported for human *Cul3* mutations associated with autism [4]. While Inc orthologs are expressed in the mammalian brain [33], their characterization in vivo remains limited. In mice, *KCTD5* deletions are homozygous lethal and cause motor deficits when heterozygous [55]; possible effects on sleep have not yet been assessed. There are no existing rodent models for mutations in *KCTD2* or *KCTD17*. Conditional genetic analysis of *inc* in flies and *Cul3* in mice underscores developmental mechanisms of action. Inc acts in developing neurons to impact sleep in adult flies [56], suggesting that its relevant substrates have developmental functions within the brain. Conditional deletion of *Cul3* in mice indicates that its behavioral phenotypes, including deficits in sociability and cognition, arise during a critical period of brain development [10], suggesting that any sleep phenotypes of *Cul3* mutants might also arise in a developmental manner. The molecular targets of Inc that influence sleep are likely conserved, as mammalian Inc orthologs restore sleep to *inc* mutants [33]. Intriguing overlaps in the synaptic phenotypes of *inc* and *Cul3* mutants in flies and mice suggest that Inc orthologs impact synaptic function through substrates that are similarly conserved. Inc and Cul3 are required in flies for synaptic homeostasis at glutamatergic neuromuscular synapses and are recruited to synapses during homeostatic potentiation [20]. *Cul3* mutations in mice alter glutamatergic neurotransmission and the abundance of glutamate receptors and other synaptic proteins [9,11]. While contributions of mammalian Inc orthologs to *Cul3* phenotypes at glutamatergic synapses remain unexplored, Cul3 and Inc orthologs are present in synaptosome preparations [33], suggesting that their targets include synaptic proteins.

Characterization of Inc orthologs has identified various substrates in cultured cells. c-Myc is a substrate for KCTD2 [57], and KCTD5 ubiquitinates ΔNp63α and RhoGDI [58,59], linking KCTD2 and KCTD5 to pathways that regulate cell proliferation. KCTD5 has also been shown to ubiquitinate a G-protein beta subunit and to regulate AKT and cAMP signaling, including in cultured striatal neurons [55,60,61]. KCTD17 ubiquitinates trichoplein, a regulator of primary cilia [62], as well as Chop and PHLPP2 [63,64], proteins that regulate lipids and adipogenesis. Whether these targets or others mediate the impact of Inc on sleep and synapses remains unknown.

Emerging evidence from genetic studies in humans suggests that Inc orthologs have functions in the brain that remain poorly understood. As noted above, *KCTD17* mutations are associated with myoclonic dystonia [37]. Polymorphisms near the *KCTD2* locus are associated with Alzheimer’s [65], and another study suggests an association of *KCTD5* and daytime sleepiness [66]. Elucidating targets of Cul3-Inc complexes within the fly brain may provide a powerful system to understand conserved mechanisms through which Cul3 and Inc contribute to nervous system function and neurological disorders.

## Acknowledgements

We thank J. Lopez for assistance with Western blots and qRT-PCR; Z. Zuchowski for assistance in constructing plasmids; A. Arzeno and A. Gonzalez for assistance with immunoprecipitations; and N. Ringstad, J. Treisman, and members of the Stavropoulos laboratory for comments on the manuscript. This work was supported by an International Student Research Fellowship from the Howard Hughes Medical Institute (HHMI) to Q.L., by a grant from the National Institutes of Health (NS091546) to D.D., and by grants from the National Institutes of Health (NS112844 and NS111304), the Mathers Foundation, Whitehall Foundation grant 2013-05-78, fellowships from the Alfred P. Sloan and Leon Levy Foundations, a NARSAD Young Investigator Award from the Brain and Behavior Foundation, the J. Christian Gillin, M.D. Research Award from the Sleep Research Society Foundation, and a Career Scientist Award from the Irma T. Hirschl/Weill-Caulier Trust to N.S.

## Author Contributions

**Conceptualization:** Q.L., D.D., N.S.

**Investigation:** Q.L., K.Y.L., R.A., F.V., X.L., W.D., J.M., and N.S.

**Resources:** Q.L., D.D., and N.S.

**Writing - original draft:** N.S. and Q.L.

**Writing - review and editing:** N.S., Q.L., and D.D.

## Materials and methods

### Plasmids and molecular cloning

#### Vectors for expression in S2 cells were as follows

pAc5.1–Inc-HA (pNS277) encodes Inc fused to a C-terminal HA epitope (<u>GSYPYDVPDYA</u>) and was generated by ligating EcoRI-XhoI digested pAc5.1/V5-HisA backbone (ThermoFisher) and an EcoRI-XhoI Inc-HA fragment from pNS273 (described below).

pAc5.1–3×Myc-Inc (pNS351) was previously described (Li et al., 2017) and encodes an N-terminal 3×Myc epitope (M<u>EQKLISEEDL</u>GS<u>EQKLISEEDL</u>GS<u>EQKLISEEDL</u>AS) fused to Inc.

pAc5.1–3×Myc-Inc^22-211^ (pNS370) was generated by ligating NheI-XhoI digested pNS309 (Stavropoulos and Young, 2011) and the NheI-XhoI fragment liberated from the PCR amplification product of pNS351 template and primers oNS315 and oNS285.

pAc5.1–3×Myc-Inc^31-211^ (pNS371) was generated similarly to pNS370, substituting primers oNS316 and oNS285.

pAc5.1–3×Myc-Inc^124-211^ (pNS372) was generated similarly to pNS370, substituting primers oNS317 and oNS285.

pAc5.1–3×Myc-Inc^1-123^ (pNS374) was generated similarly to pNS370, substituting primers oNS277 and oNS319.

pAc5.1–3×Myc-Inc^1-156^ (pNS375) was generated similarly to pNS370, substituting primers oNS277 and oNS320.

pAc5.1–3×Myc-Inc^1-186^ (pNS376) was generated similarly to pNS370, substituting primers oNS277 and oNS321.

pAc5.1–3×Myc-Inc^22-123^ (pNS377) was generated similarly to pNS370, substituting primers oNS315 and oNS319.

pAc5.1–3×HA-Inc (pNS402) encodes an N-terminal 3×HA epitope (<u>MYPYDVPDYA</u>GS<u>YPYDVPDYA</u>GS<u>YPYDVPDYAAS</u>) fused to Inc and was generated by ligating NheI-XhoI digested pNS310 (Stavropoulos and Young, 2011) and an NheI-XhoI *inc* fragment liberated from pNS351.

pAc5.1–3×FLAG-Cul3 (pNS403) encodes an N-terminal 3×FLAG epitope (M<u>DYKDDDDK</u>GS<u>DYKDDDDK</u>GS<u>DYKDDDDK</u>AS) fused to *Drosophila* Cul3 and was generated by ligating NheI-NotI digested pNS311 and an NheI-NotI Cul3 fragment liberated from pNS314 (Stavropoulos and Young, 2011). pNS311 contains an N-terminal 3×FLAG tag and was generated from pNS298, a pAc5.1/V5-HisA derivative that contains a C-terminal 3×FLAG tag. To construct pNS298, oligonucleotides oNS234 and oNS235 were phosphorylated, annealed, and cloned into XhoI-XbaI digested pAc5.1/V5-HisA. To construct pNS311, EcoRI-NotI digested pAc5.1/V5-HisA was ligated to the EcoRI-NotI fragment liberated from the PCR amplification product of pNS298 template and primers oNS240 and oNS241.

pAc5.1–3×FLAG-Inc (pNS408) encodes an N-terminal 3×FLAG epitope (<u>MDYKDDDDK</u>GS<u>DYKDDDDK</u>GS<u>DYKDDDDKAS</u>) fused to Inc and was generated by ligating NheI-NotI digested pNS311 and an NheI-XhoI *inc* fragment liberated from pNS351.

pAc5.1–3×HA-Inc^F47A^ (pNS409) encodes an N-terminal 3×HA epitope (M<u>YPYDVPDYA</u>GS<u>YPYDVPDYA</u>GS<u>YPYDVPDYA</u>AS) fused to Inc^F47A^ and was generated by ligating NheI-XhoI digested pNS310 and an NheI-XhoI digested Inc^F47A^ fragment. The Inc^F47A^ fragment was prepared by fusion PCR in two steps. First, overlapping 5’ and 3’ Inc^F47A^ fragments were generated by PCR amplification of pNS408 template with primers oNS277/oNS695 and oNS684/oNS694, respectively. An equimolar mix of these fragments was then used as template for fusion PCR amplification with primers oNS277 and oNS684.

pAc5.1–3×HA-Inc^R50E^ (pNS410) was generated similarly to pNS409, substituting primers oNS277/oNS697 and oNS684/oNS696 to obtain 5’ and 3’ Inc^R50E^ fragments, respectively.

pAc5.1–3×HA-Inc^D57A^ (pNS411) was generated similarly to pNS409, substituting primers oNS277/oNS699 and oNS684/oNS698 to obtain 5’ and 3’ Inc^D57A^ fragments, respectively.

pAc5.1–3×HA-Inc^D61A^ (pNS412) was generated similarly to pNS409, substituting primers oNS277/oNS701 and oNS684/oNS700 to obtain 5’ and 3’ Inc^D61A^ fragments, respectively.

pAc5.1–3×HA-Inc^E104K^ (pNS413) was generated similarly to pNS409, substituting primers oNS277/oNS703 and oNS684/oNS702 to obtain 5’ and 3’ Inc^E104K^ fragments, respectively.

pAc5.1–3×HA-Inc^F105A^ (pNS419) was generated similarly to pNS409, substituting primers oNS277/oNS1123 and oNS684/oNS1122 to obtain 5’ and 3’ Inc^F105A^ fragments, respectively.

pAc5.1–3×HA-Inc^Y106F^ (pNS420) was generated similarly to pNS409, substituting primers oNS277/oNS1146 and oNS684/oNS1145 to obtain 5’ and 3’ Inc^Y106F^ fragments, respectively.

pAc5.1–3×HA-Inc^N107A^ (pNS421) was generated similarly to pNS409, substituting primers oNS277/oNS1125 and oNS684/oNS1124 to obtain 5’ and 3’ Inc^N107A^ fragments, respectively.

pAc5.1–3×FLAG-Inc^D73A^ (pNS430) and pAc5.1–3×FLAG-Inc^N82A^ (pNS431) were generated simultaneously, using PCR amplification of pNS408 template with primers oNS277/oNS686 and oNS684/oNS685 to generate overlapping Inc^N82A^ and Inc^D73A^ fragments, respectively. An equimolar mix of these fragments was used as template for fusion PCR amplification with primers oNS277 and oNS684, and the PCR product was digested with NheI-XhoI and ligated to NheI-XhoI digested pNS311. Sequencing of multiple clones identified both single mutants.

pAc5.1–3×FLAG-Inc^K88D^ (pNS432) and pAc5.1–3×FLAG-Inc^E101K^ (pNS433) were generated similarly to pNS430 and pNS431, substituting primers oNS277/oNS688 and oNS684/oNS687 to obtain overlapping Inc^E101K^ and Inc^K88D^ fragments, respectively.

pAc5.1–3×FLAG-Inc^T36A^ (pNS434) and pAc5.1–3×FLAG-Inc^D71A^ (pNS435) were generated similarly to pNS430 and pNS431, substituting primers oNS277/oNS693 and oNS684/oNS689 to obtain overlapping Inc^D71A^ and Inc^T36A^ fragments, respectively.

pAc5.1–3×HA-Inc^R85E^ (pNS436) was generated similarly to pNS430 and pNS431, substituting primers oNS277/oNS692 and oNS684/oNS691 to obtain overlapping Inc^R85E^ and Inc^D71A^ fragments, respectively, and NheI-XhoI digested pNS310. Inc^R85E^ was identified by sequencing multiple clones.

pAc5.1–3×HA-Inc^F47A/F105A^ (pNS422) was generated similarly to pNS409, substituting pNS409 as template and primers oNS277/oNS1123 and oNS684/oNS1122 to obtain 5’ and 3’ Inc^F47A/F105A^ fragments, respectively.

pAc5.1–3×HA-Inc^D73A/N82A^ (pNS414) was generated similarly to pNS409, substituting pNS430 template and primers oNS277/oNS686 to obtain the 5’ Inc^D73A/N82A^ fragment and pNS431 template and primers oNS684/oNS685 to obtain the 3’ Inc^D73A/N82A^ fragment.

pAc5.1–3×HA-Inc^K88D/E101K^ (pNS415) was generated similarly to pNS409, substituting pNS432 template and primers oNS277/oNS688 to obtain the 5’ Inc^K88D/E101K^ fragment and pNS433 template and primers oNS684/oNS687 to obtain the 3’ Inc^K88D/E101K^ fragment.

pAc5.1–3×HA-Inc^T36A/D71A^ (pNS416) was generated similarly to pNS409, substituting pNS434 template and primers oNS277/oNS693 to obtain the 5’ Inc^T36A/D71A^ fragment and pNS435 template and primers oNS684/oNS689 to obtain the 3’ Inc^T36A/D71A^ fragment.

pAc5.1–3×HA-Inc^D71A/R85E^ (pNS417) was generated similarly to pNS409, substituting pNS435 template and primers oNS277/oNS692 to obtain the 5’ Inc^D71A/R85E^ fragment and pNS436 template and primers oNS684/oNS691 to obtain the 3’ Inc ^D71A/R85E^ fragment.

pAc5.1–3×HA-Inc^T36A/D71A/R85E^ (pNS418) was generated by ligating EcoRV-XhoI digested pNS434 backbone and an Inc^D71A/R85E^ fragment obtained by EcoRV-XhoI digestion of the PCR amplification product of pNS417 template and primers oNS277 and oNS684.

pAc5.1–3×HA-Inc^R135H^ (pNS426) was generated by ligating HindIII-XhoI digested pNS402 backbone and a HindIII-XhoI Inc^R135H^ fragment. The Inc^R135H^ fragment was prepared by fusion PCR in two steps. First, overlapping 5’ and 3’ Inc^R135H^ fragments were generated by PCR amplification of pNS346 template with primers oNS1126/oNS1555 and oNS1127/oNS1554, respectively. An equimolar mix of these fragments was then used as template for fusion PCR amplification with primers oNS1126 and oNS1127. pUASTattB–Myc-Inc (pNS346) encodes an N-terminal Myc epitope (M<u>EQKLISEEDL</u>AS) fused to Inc, as previously described (Li et al., 2017).

pAc5.1–3×HA-Inc^AAA^ (pNS427) was generated similarly to pNS426, substituting a HindIII-XhoI Inc^AAA^ fragment generated similarly with fusion PCR; primer pairs oNS1126/oNS1557 and oNS1127/oNS1556 were used to obtain 5’ and 3’ Inc^AAA^ fragments, respectively.

pAc5.1–3×Myc-Inc^T36A/D71A/R85E^ (pNS437) was generated by ligating NheI-XhoI digested pNS309 backbone and an NheI-XhoI Inc^T36A/D71A/R85E^ fragment liberated from pNS418.

pAc5.1–3×Myc-Inc^F47A^ (pNS438) was generated by ligating NheI-XhoI digested pNS309 backbone and an NheI-XhoI Inc^F47A^ fragment liberated from pNS409.

pAc5.1–3×Myc-Inc^F105A^ (pNS439) was generated by ligating NheI-XhoI digested pNS309 backbone and an NheI-XhoI Inc^F105A^ fragment liberated from p419.

pAc5.1–3×Myc-Inc^F47A/F105A^ (pNS440) was generated by ligating NheI-XhoI digested pNS309 backbone and an NheI-XhoI Inc^F47A/F105A^ fragment liberated from pNS422.

pAc5.1–3×Myc-Inc^R135H^ (pNS441) was generated by ligating HindIII-XhoI digested pNS351 backbone and a HindIII-XhoI Inc^R135H^ fragment prepared as for pNS426.

pAc5.1–3×Myc-Inc^AAA^ (pNS442) was generated by ligating HindIII-XhoI digested pNS351 backbone and a HindIII-XhoI Inc^AAA^ fragment prepared as for pNS427.

#### Vectors for *Drosophila* transgenesis were as follows

pUASTattB–Myc-Inc^R135H^ (pNS428) was generated by ligating a BglII-XhoI digested pUASTattB backbone (Bischof et al., 2007), prepared by liberating an unrelated BglII-XhoI insert, and a BgII-XhoI digested Myc-Inc^R135H^ fragment, generated with fusion PCR as for pNS426.

pUASTattB–Myc-Inc^AAA^ (pNS429) was generated by ligating a BgII-XhoI pUASTattB backbone prepared from pNS349 [33] and a BgII-XhoI digested Myc-Inc^AAA^ fragment, generated with fusion PCR as for pNS427.

pUAST–Inc-HA (pNS273) was previously described and encodes Inc fused to a C-terminal HA epitope (GS<u>YPYDVPDYA</u>) [33].

pUASTattB–3×FLAG-Inc (pNS404) encodes an N-terminal 3×FLAG epitope (<u>MDYKDDDDK</u>GS<u>DYKDDDDK</u>GS<u>DYKDDDDK</u>AS) fused to Inc and was generated by three-piece ligation of EcoRI-XhoI digested pUASTattB, an EcoRI-NheI 3×FLAG fragment liberated from the PCR amplification product of pNS311 template and primers ACF and oNS241, and an NheI-XhoI *inc* fragment liberated from pNS351.

pUASTattB–3×FLAG-Inc^1-186^ (pNS405) was generated similarly to pNS404, substituting an NheI-XhoI Inc^1-186^ fragment liberated from pNS376.

pUASTattB–3×FLAG-Inc^1-156^ (pNS406) was generated similarly to pNS404, substituting an NheI-XhoI Inc^1-156^ fragment liberated from pNS375.

pUASTattB–3×FLAG-Inc^1-123^ (pNS407) was generated similarly to pNS404, substituting an NheI-XhoI Inc^1-123^ fragment liberated from pNS374.

pUASTattB–3×FLAG-Inc^F47A^ (pNS423) was generated similarly to pNS404, substituting an NheI-XhoI Inc^F47A^ fragment liberated from pNS409.

pUASTattB–3×FLAG-Inc^F105A^ (pNS424) was generated similarly to pNS404, substituting an NheI-XhoI Inc^F105A^ fragment liberated from pNS419.

pUASTattB–3×FLAG-Inc^F47A/F105A^ (pNS425) was generated similarly to pNS404, substituting an NheI-XhoI Inc^F47A/F105A^ fragment liberated from pNS422.

### Oligonucleotides

#### Oligonucleotides used in this work, listed 5’ to 3’, are as follows

oNS180 ACGTAGATCTGAACCGCAGCAGCGGCAACACCATC

oNS184 CCAGCCATCCGACAGCGTTGAGATC

oNS234 TCGAGGCTAGCGACTACAAGGATGATGACGATAAGGGCTCCGATTACAAGGACGACGATGATAAGGGATCCGATTACAAGGATGATGACGACAAGTGAT

oNS235 CTAGATCACTTGTCGTCATCATCCTTGTAATCGGATCCCTTATCATCGTCGTCCTTGTAATCGGAGCCCTTATCGTCATCATCCTTGTAGTCGCTAGCC

oNS240 ACTGGAATTCCGCGGCAACATGGACTACAAGGATGATGACGATAAGGGC

oNS241 ACTGGCGGCCGCTCCTAGGGTGCTAGCCTTGTCGTCATCATCCTTGTAATCGGAT

oNS277 ACGTGCTAGCATGAGCACGGTGTTCATAAACTCGC

oNS285 ACGTGCTAGCTCGAGGGGTTGTGTGTGAATATATAGCGCGA

oNS315 ACGTGCTAGCCAGTGGGTCAAGCTGAACGTAG

oNS316 ACGTGCTAGCACCTACTTCCTCACCACAAAGACG

oNS317 ACGTGCTAGCCAGCGACCCCAAACGGACAA

oNS319 ACGTCTCGAGTTAATCCCTGTGCAGGATGCACTC

oNS320 ACGTCTCGAGTTACCTCCAGCCATCCGACAG

oNS321 ACGTCTCGAGTTATGTGCCACACTCTTTGGATACCA

oNS337 TTTTTTTTTTTTTTTTTTTTTTTVN

oNS684 ACGTCTCGAGGGGTTGTGTGTGAATATATAGCGCGA

oNS685 CCTGATCGACAGAGCCCCCAAATACTTTGC

oNS686 GGCGCAGGTAGGCGAGCACGGGTGCAAA

oNS687 TGCGCCACGGCGACCTTGTGCTCGAT

oNS688 TAGAACTCAGCCTCCTTCAGGACGCCTTC

oNS689 CTACTTCCTCACCGCCAAGACGACGCTC

oNS691 GCCTACCTGATCGCCAGAGACCCCAAA

oNS692 CAAGCTTGCCGTGTTCCAGGTAATTGAGC

oNS693 TTTGGGGTCTCTGGCGATCAGGTAGGC

oNS694 GACCCAAATTCGGCCCTCTCCCGTCTG

oNS695 CAGACGGGAGAGGGCCGAATTTGGGTC

oNS696 TCGTTCCTCTCCGAACTGATTCAGGAGG

oNS697 CCTCCTGAATCAGTTCGGAGAGGAACGA

oNS698 CAGGAGGACTGCGCCTTGATATCAGATCG

oNS699 CGATCTGATATCAAGGCGCAGTCCTCCTG

oNS700 CGACTTGATATCAGCCCGGGACGAGAC

oNS701 GTCTCGTCCCGGGCTGATATCAAGTCG

oNS702 CCTGGAGGAGGCTAAGTTCTACAACGTGAC

oNS703 GTCACGTTGTAGAACTTAGCCTCCTCCAGG

oNS1122 GGAGGCTGAGGCCTACAACGTGAC

oNS1123 GTCACGTTGTAGGCCTCAGCCTCC

oNS1124 GCTGAGTTCTACGCCGTGACGCAGC

oNS1125 GCTGCGTCACGGCGTAGAACTCAGC

oNS1126 CAACTGCAACTACTGAAATCTGCCAAGAAG

oNS1127 GGTAGTTTGTCCAATTATGTCACACCACAGAAG

oNS1145 GAGGCTGAGTTCTTCAACGTGACGCAGC

oNS1146 GCTGCGTCACGTTGAAGAACTCAGCCTC

oNS1554 AAGCGCGTTTATCATGTGCTGCAGTGC

oNS1555 GCACTGCAGCACATGATAAACGCGCTT

oNS1556 ATCAGCATGCAGTACACGGCCGCCGCGCCCTTCGAAAACAATGAGTTCCTG

oNS1557 ATTGTTTTCGAAGGGCGCGGCGGCCGTGTACTGCATGCTGATCAGCT

oNS1783 TGTCCACGTACCAGATGTGTGTGCTA

oNS1784 ACTAGCAAGTTGTGAGCCAGACGC

ACF GACACAAAGCCGCTCCATCAG

RPS3A CGAACCTTCCGATTTCCAAGAAACGC

RPS3B ACGACGGACGGCCAGTCCTCC

### Cell culture and biochemistry

S2 cells were cultured in S2 media containing 10% FBS, penicillin, and streptomycin, and were transfected with Effectene (Qiagen) as described previously (Stavropoulos and Young, 2011). Transfections were performed in 6 well plates for ∼24 hr, after which liposome-containing media was replaced with fresh culture media. 400 ng of total DNA was used for each transfection. For transfections involving two plasmids, an equal amount of each was used. For comparisons of multiple plasmids, empty vector lacking insert was used to equalize DNA amounts as required. Cells were harvested 36-48 hr after transfection, washed in PBS, and lysed in ice-cold NP40 lysis buffer (50 mM Tris pH 7.6, 150mM NaCl, 0.5% NP40) containing protease inhibitors (Sigma, P8340). Protein extracts were quantitated in duplicate (BioRad, 5000111). For measurements of protein stability, 100 μg/ml cycloheximide (Fisher, AC357420010) was added at specified timepoints prior to lysis.

For immunoprecipitations of truncated Inc proteins and Inc point mutants from S2 cells, 300-1000 µg of total protein was incubated overnight at 4**°**C with 1:100 anti-FLAG (Sigma, F1804), anti-Myc (Sigma, C3956), or rat anti-Inc antibody [17], followed by incubation with 20 µl (50% slurry) of protein G sepharose beads (ThermoFisher, 10-1243) at 4**°**C on a nutator for 1 hr or overnight. Alternatively, 20 µl (50% slurry) of anti-FLAG (Sigma, F2426) or anti-Myc affinity gel (Sigma, E6654) was used to immunoprecipitate samples. Samples were washed 4×5 min at 4**°**C with NP40 lysis buffer, denatured in SDS sample buffer, separated on Tris SDS-PAGE gels, and transferred to nitrocellulose.

Protein extracts were prepared from whole flies or sieved heads by manual pestle homogenization in ice-cold NP40 lysis buffer containing protease inhibitors. To assess in vivo expression of Inc proteins, 30 µg of lysate was separated on Tris-SDS-PAGE gels. For immunoprecipitations of whole fly extracts, 800 µg of protein was incubated with 30 µl (50% slurry) of anti-Myc affinity gel for 1.5 hr at 4**°**C on a nutator. Samples were washed, denatured, electrophoresed, and transferred to membranes as described above.

### Western blots and antibodies

Membranes were incubated for 60-90 min at room temperature in blocking buffer (LI-COR, 927-40000). Membranes were subsequently incubated for 1-2 hr at room temperature or 4**°**C overnight in blocking buffer containing 0.1% Tween 20 and the appropriate primary antibodies: rabbit anti-Myc (1:2,000, Sigma, C3956), mouse anti-FLAG (1:2,000, Sigma, F1804), rat anti-HA (1:2,000, Roche, 11867431001), and rabbit anti-HA (1:2,000, Bethyl Laboratories, A190-208A), mouse anti-tubulin (1:10,000, DSHB, 12G10), and rabbit anti-tubulin (1:60,000, VWR, 89364-004). Membranes were washed 4×5 min in TBST (150 mM NaCl, 10mM Tris pH 7.6, and 0.1% Tween 20) and subsequently incubated in the dark for 30-60 min at room temperature in blocking buffer containing 0.1% Tween 20, 0.01% SDS, and appropriate secondary antibodies, all diluted 1:15,000 or 1:30,000: Alexa 680 donkey anti-rabbit (Life Technologies, A10043), Alexa 790 anti-rabbit (Life Technologies, A11374), Alexa 680 donkey anti-mouse (Life Technologies, A10038), Alexa 790 anti-mouse (Life Technologies, A11371), Alexa 790 anti-rat (Jackson ImmunoResearch, 712-655-153), and Alexa 680 donkey anti-rat (Jackson ImmunoResearch, 712-625-1533). Membranes were then washed 4×5 min in TBST, 1×5 min in TBS, and imaged on a Li-Cor Odyssey CLx instrument.

### qRT-PCR

Total RNA was isolated from adult male fly heads with TRIZOL (ThermoFisher, 15596-026). 5 µg of RNA was reverse transcribed with d(T)_23_VN primer (oNS337) and SuperScript II reverse transcriptase (ThermoFisher, 18064014). qPCR was performed using a BioRad DNA Engine Opticon 2 System, SYBR Green Supermix (Bio-Rad, 1725272), and the following primers: oNS180 and oNS184 (*inc*); oNS1783 and oNS1784 (*Cul3*); RPS3A and RPS3B (*rps3*).

### Fly stocks and transgenes

*elav^c155^-Gal4* [67], *inc^1^* [17], *inc^kk3^* [20], *inc-Gal4* [17], *inc^1^ inc-Gal4* [33], *BG57-Gal4* [68], *attP2: UAS-Myc-Inc* [33], *attP2: UAS-3×FLAG-Inc* [33], and *UAS-3×FLAG-3×HA-Cul3* [69] were previously described. *UAS-Cul3-RNAi* (11861R-2) and *UAS-inc-RNAi* (18225) were obtained from the NIG-Fly and VDRC stock centers, respectively. Transgenic flies generated in this study using pUASTattB-based vectors were integrated at *attP2* [70] with phiC31 recombinase (BestGene). All transgenes were backcrossed six to eight generations to Bloomington stock 5905, an isogenic *w^1118^* stock described elsewhere as iso31 [71].

### Sleep analysis

Crosses were set with five virgin females and three males on cornmeal, agar, and molasses food. One- to four-day old male flies eclosing from LD-entrained cultures raised at 25**°**C were loaded in glass tubes containing cornmeal, agar, and molasses food. Animals were monitored for 5-7 days at 25**°**C in LD cycles using DAM2 monitors (Trikinetics). The first 36-48 hours of data were discarded and an integral number of days of data (3-5) were analyzed using custom Matlab code. Locomotor data were collected in 1 min bins. Sleep was defined by locomotor inactivity for 5 min or more; all minutes within inactive periods exceeding 5 min were assigned as sleep. Dead animals were excluded from analysis by a combination of automated filtering and visual inspection of locomotor traces. Two to four independent biological replicates were performed for all sleep assays.

### Electrophysiology

Two-electrode voltage clamp (TEVC) recordings were performed in muscle 6 in abdominal segments 2 and 3, as previously described [20]. Muscles were clamped at −70 mV, with a leak current below 5 nA. mEPSCs were recorded for 1 min from each muscle cell in the absence of stimulation. Twenty EPSCs were acquired for each cell under stimulation at 0.5 Hz, using 0.5 ms stimulus duration and with stimulus intensity adjusted with an ISO-Flex Stimulus Isolator (A.M.P.I.). To acutely block postsynaptic receptors, larvae were incubated with philanthotoxin-433 (PhTx; 20 μM; Sigma) in HL-3 for 10 min. Data were analyzed using Clampfit 10.7 (Molecular Devices), MiniAnalysis (Synaptosoft), Excel (Microsoft), and GraphPad Prism (GraphPad Software).

### Statistics

Normality of data and equal variance were assessed with Shapiro-Wilk and Brown-Forsythe tests, respectively. Kruskal-Wallis and Dunn’s post-hoc tests were used for comparisons of total sleep, daytime sleep, nighttime sleep, sleep bout length, sleep bout number, and electrophysiological parameters. For other data, one-way ANOVA and Tukey post-hoc tests were used for data that met the assumption of normality and equal variance; for normally distributed data with unequal variance, Welch’s ANOVA and Dunnet’s post-hoc tests were used. Otherwise, Kruskal-Wallis and Dunn’s post-hoc tests were used.

### Sequence alignments

Alignments were performed with Clustal Omega 2.1 and BOXSHADE. GenBank accession numbers for proteins in Figure S1A are: Inc, NP_001284787; KCTD2, NP_056168; KCTD5, NP_061865; KCTD17.4, NP_001269615.

## Supporting Information

**S1 Fig.**
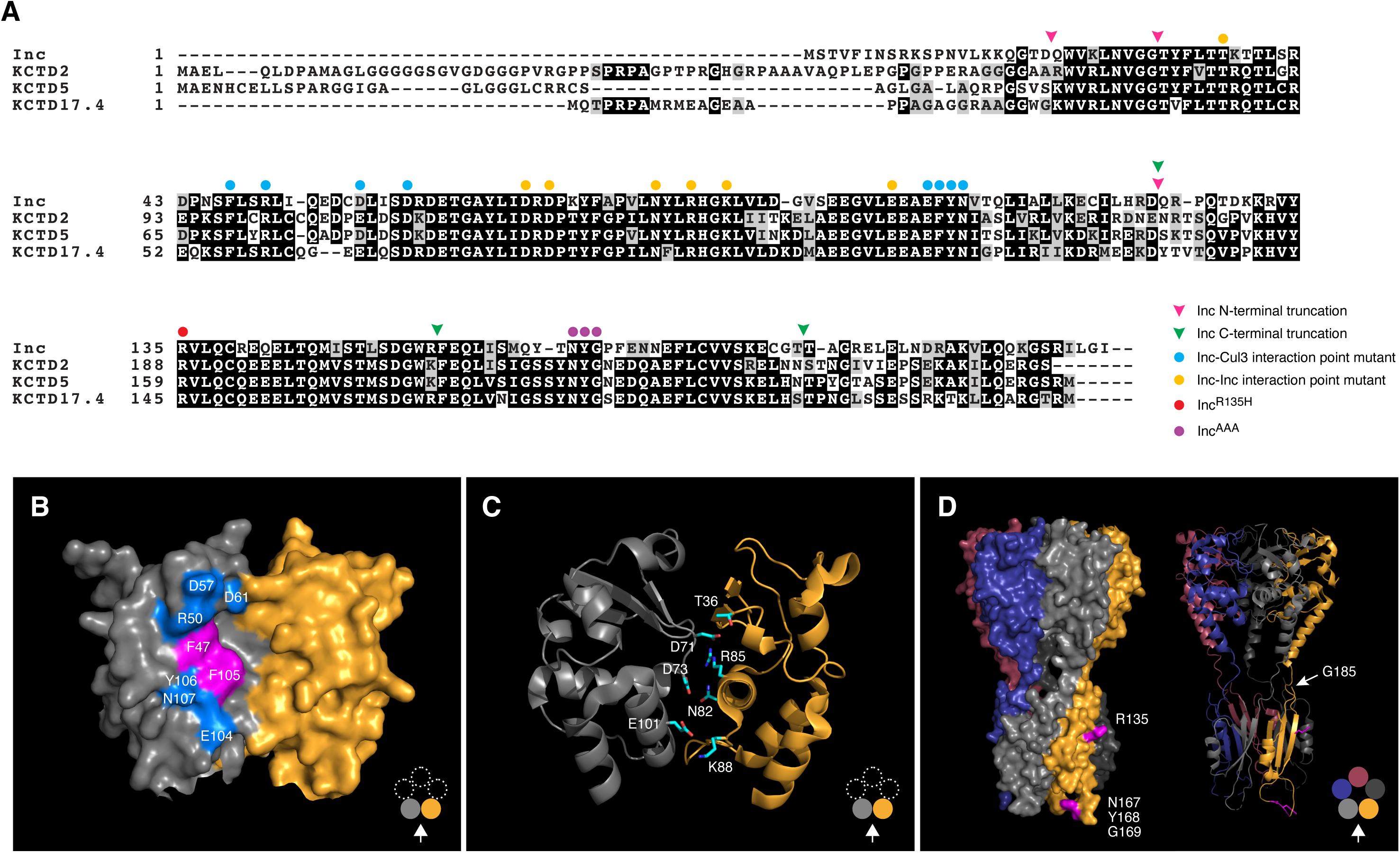
Mutagenesis of Inc. **(A)** Alignment of Inc and its human orthologs. Identical and similar residues are shaded in black and gray respectively. Locations of Inc truncations and point mutants are indicated by arrowheads and closed circles, respectively. **(B)** Surface rendering of adjacent BTB domains from the crystal structure of human KCTD5 [39]. Here and in subsequent figure panels, Inc residue numbers are superimposed on conserved equivalents in KCTD5. Inc residues whose mutation specifically weakens Inc-Cul3 associations are highlighted in magenta; other mutated residues are highlighted in blue. **(C)** Ribbon rendering of adjacent KCTD5 BTB domains. Side chains are shown for Inc residues mutated to assess Inc multimerization. Note that the T36A/D71A/R85E triple mutant alters both faces of Inc. **(D)** Surface (left) and ribbon (right) rendering of human KCTD5. Inc C-terminal point mutants are labeled and indicated in magenta; side chains are shown in the ribbon rendering. Note that KCTD5 residues equivalent to Inc amino acids 186-211 are not resolved in the crystal structure, suggesting that they are disordered.

**S2 Fig.**
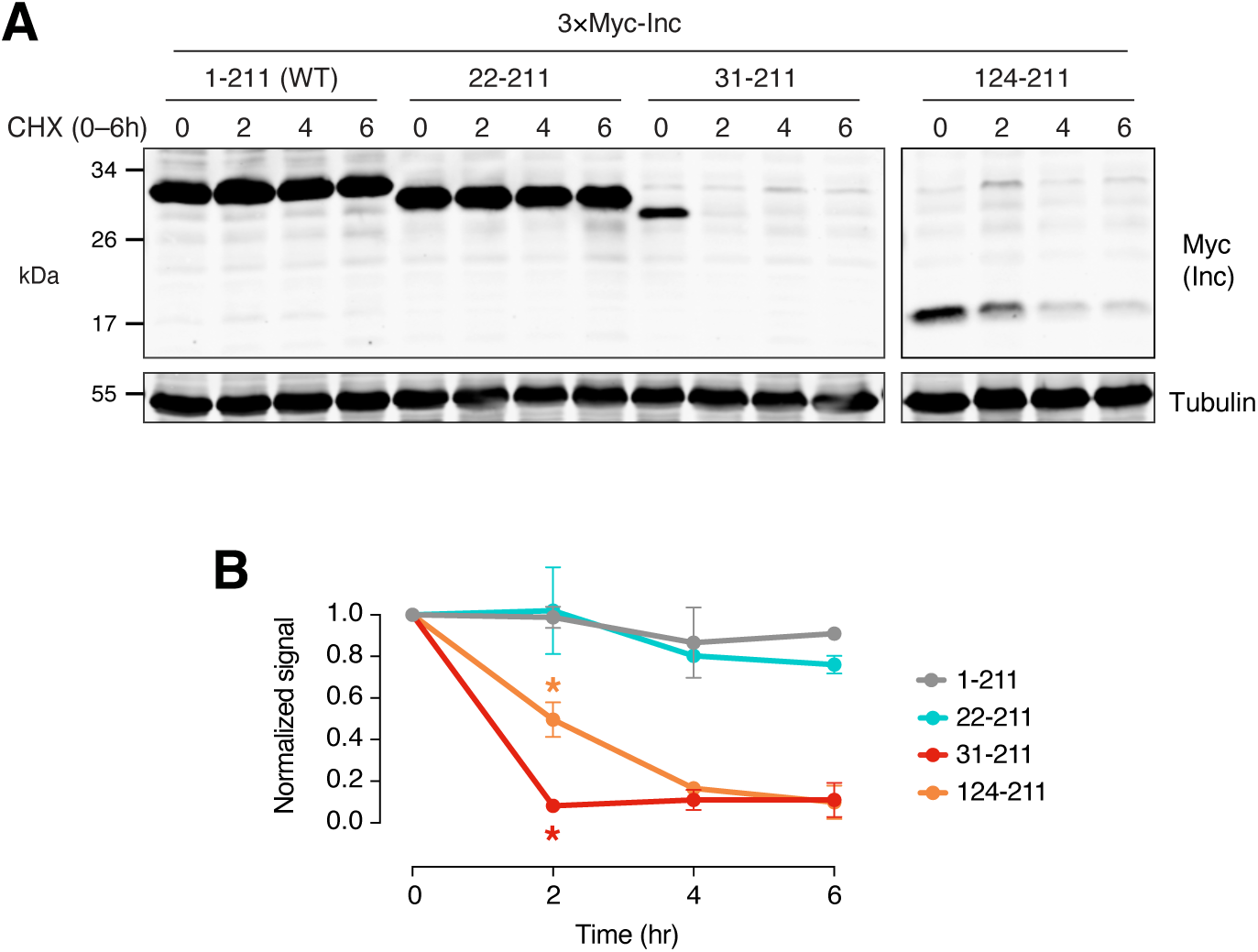
Stability of Inc truncations in S2 cells. **(A)** Immunoblot of lysates prepared from transiently transfected S2 cells treated with cycloheximide (CHX) for indicated durations. **(B)** Quantitation of two biological replicates. For all timepoints, Welch’s ANOVA p<0.05 for comparison to wild-type Inc (1-211); Dunnett’s tests; *p < 0.05.

**S3 Fig.**
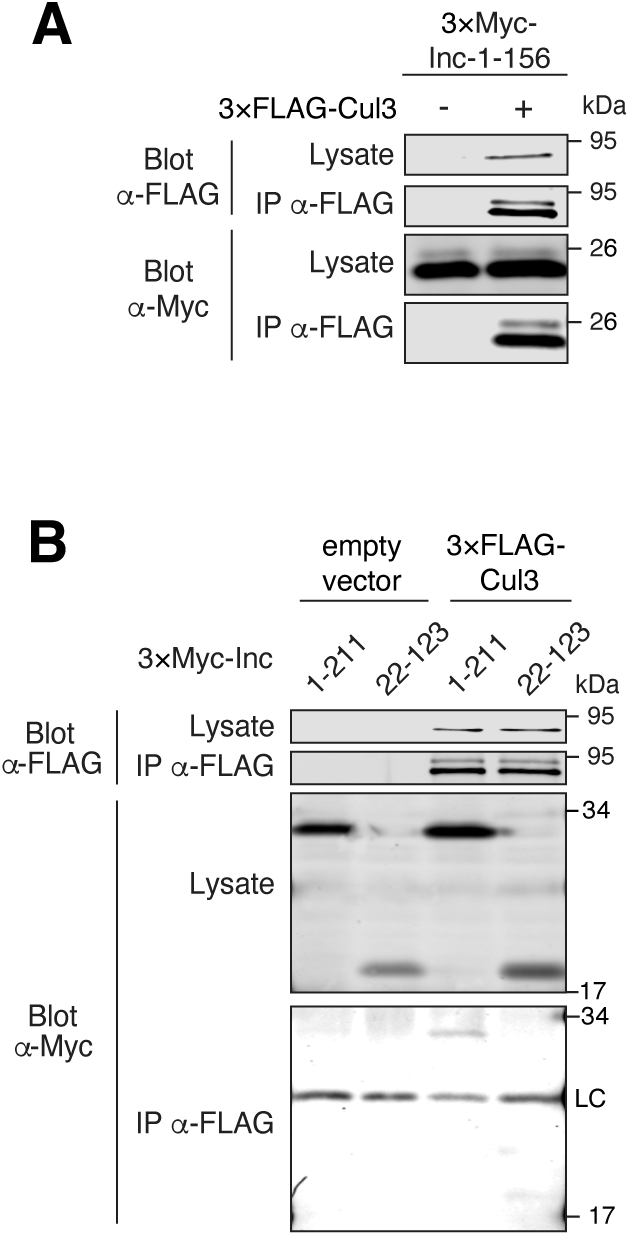
Interactions of Inc 1-156 and Inc 22-123 with Cul3. **(A-B)** Co-immunoprecipitation of 3×Myc-tagged Inc 1-156 **(A)** or Inc 22-123 **(B)** with 3×FLAG-Cul3 from transiently transfected S2 cells.

**S4 Fig.**
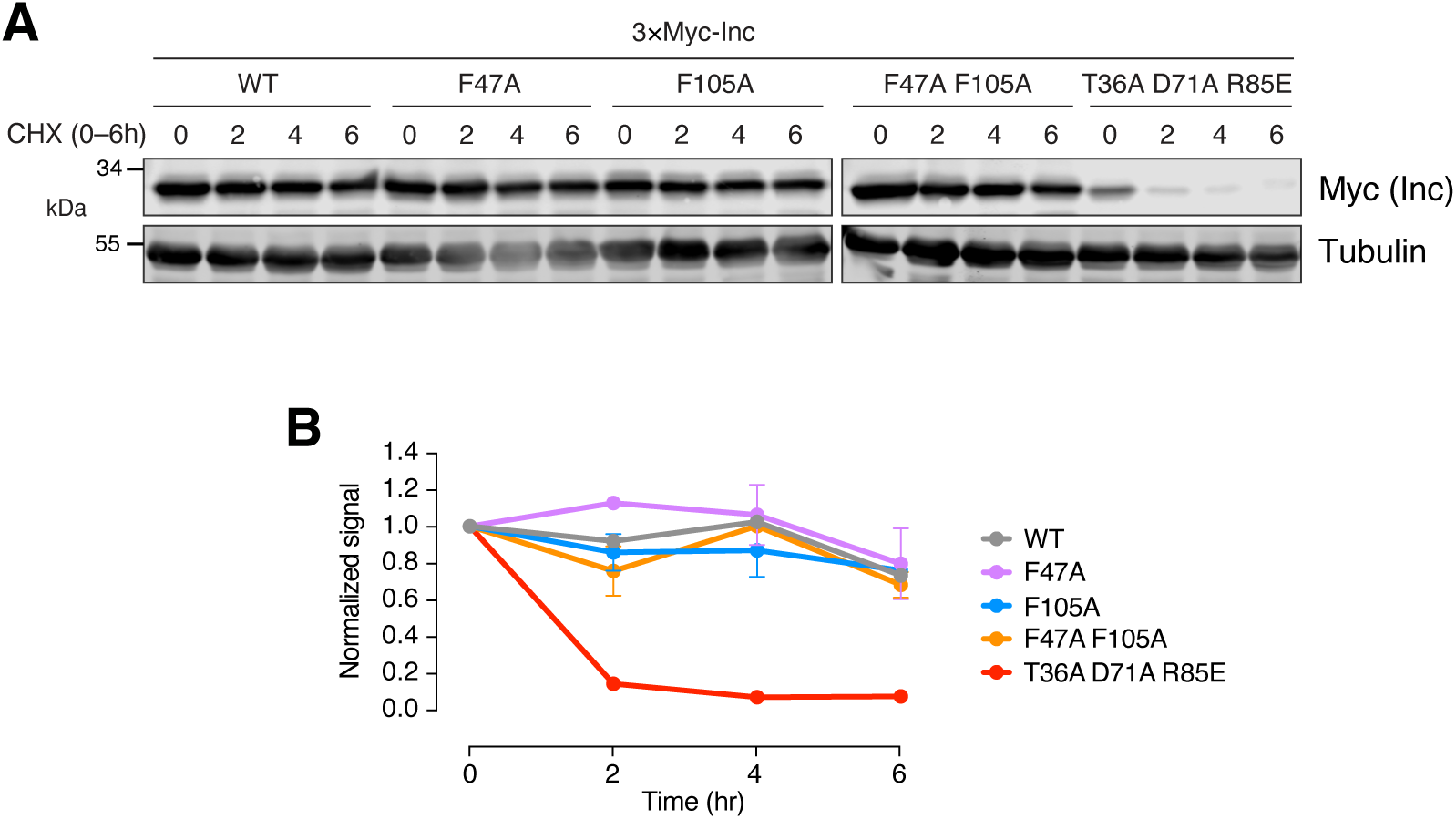
Stability of Inc point mutants in S2 cells. **(A)** Immunoblot of lysates prepared from transiently transfected S2 cells treated with cycloheximide (CHX) for indicated durations. **(B)** Quantitation of two biological replicates. For all timepoints, Welch’s ANOVA p < 0.05 for comparison to wild-type Inc.

**S5 Fig.**
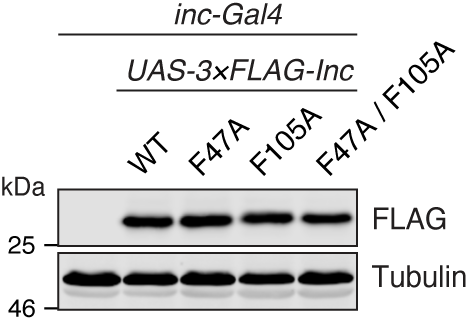
Expression of Inc BTB domain point mutants in vivo. Immunoblot of male whole animal lysates expressing 3×FLAG-tagged Inc and Inc point mutants under *inc-Gal4* control. Animals are heterozygous for UAS transgenes.

**S6 Fig.**
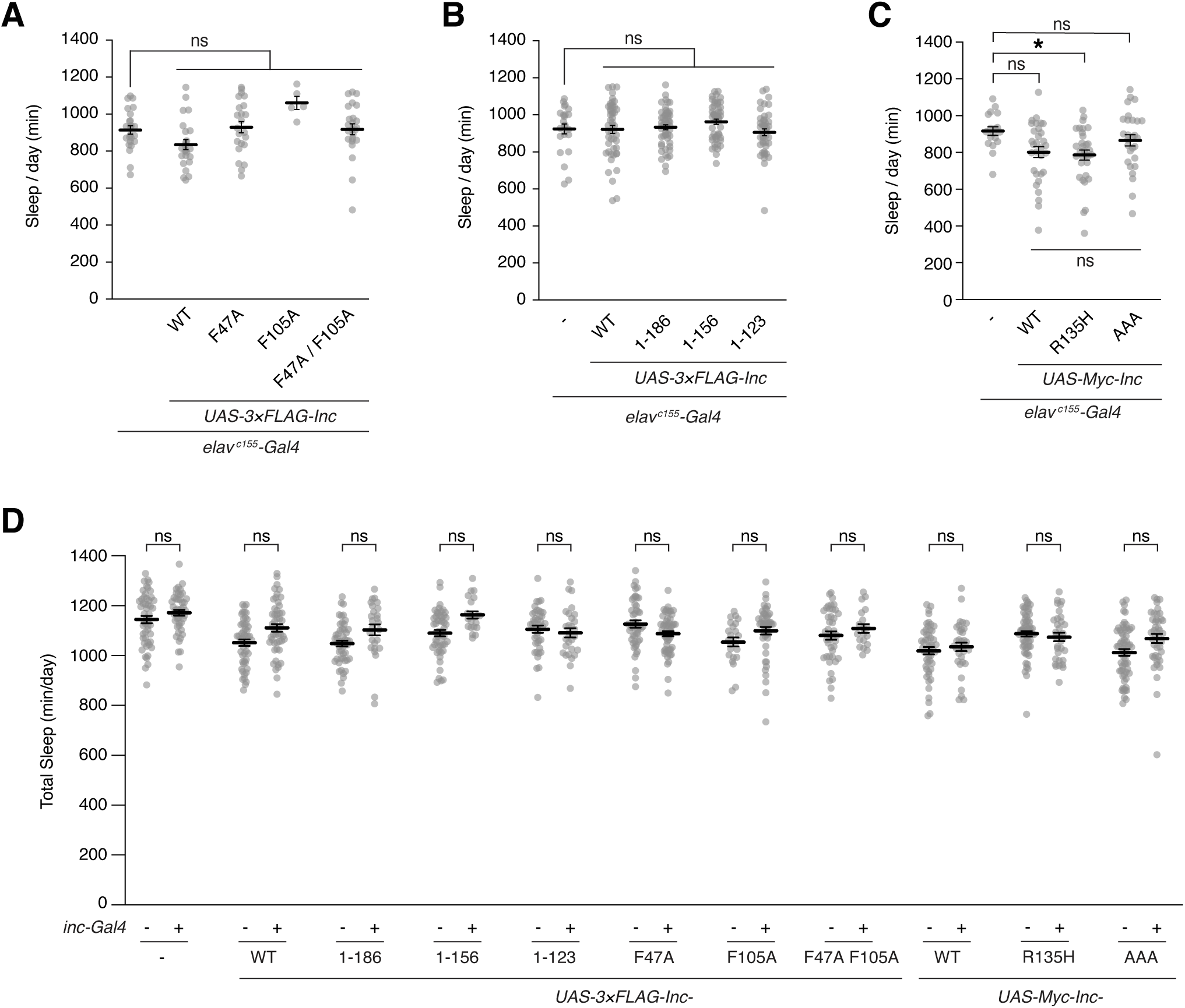
Inc truncations and point mutants lack dominant negative activity in vivo. Total daily sleep for animals expressing tagged Inc or Inc point mutants under the control of *elav^c155^-Gal4* **(A-C)** and *inc-Gal4* **(D)**. **(A)** *elav^c155^-Gal4* expression of 3×FLAG-tagged Inc point mutants. n = 5-24. **(B)** *elav^c155^-Gal4* expression of 3×FLAG-tagged Inc truncations. n = 23-54. **(C)** *elav^c155^-Gal4* expression of Myc-tagged Inc point mutants. Note that Inc^R135H^ is statistically indistinguishable from Inc and Inc^AAA^. n = 18-32. **(D)** *inc-Gal4* expression of 3×FLAG-tagged Myc-tagged Inc point mutants. n = 19-63; For all panels, mean ± SEM is shown. Kruskal-Wallis and Dunn’s tests; *p < 0.01; ns, not significant (p > 0.05). For all panels, animals are heterozygous for UAS transgenes.

**S7 Fig.**
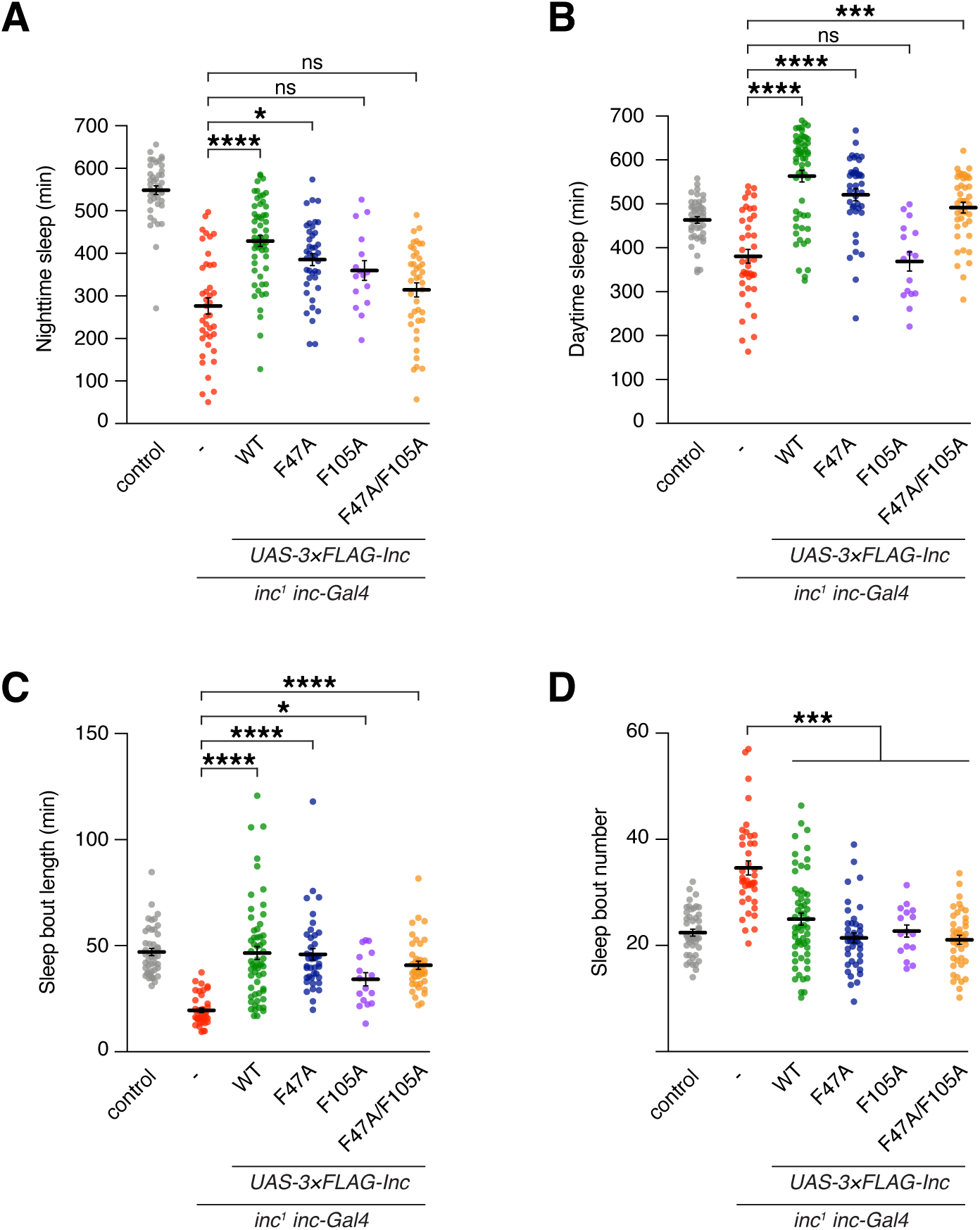
Additional sleep parameters for animals expressing Inc BTB domain point mutants. **(A-D)** Sleep parameters for *inc^1^ inc-Gal4* animals expressing 3×FLAG-tagged Inc or Inc point mutants. **(A)** Nighttime sleep. **(B)** Daytime sleep. **(C)** Sleep bout duration. **(D)** Sleep bout number. Mean ± SEM is shown. n = 16-58 as in Fig 3B; Kruskal-Wallis and Dunn’s tests; *p < 0.05; ***p < 0.001; ****p < 0.0001; ns, not significant (p > 0.05). For all panels, animals are heterozygous for UAS transgenes.

**S8 Fig.**
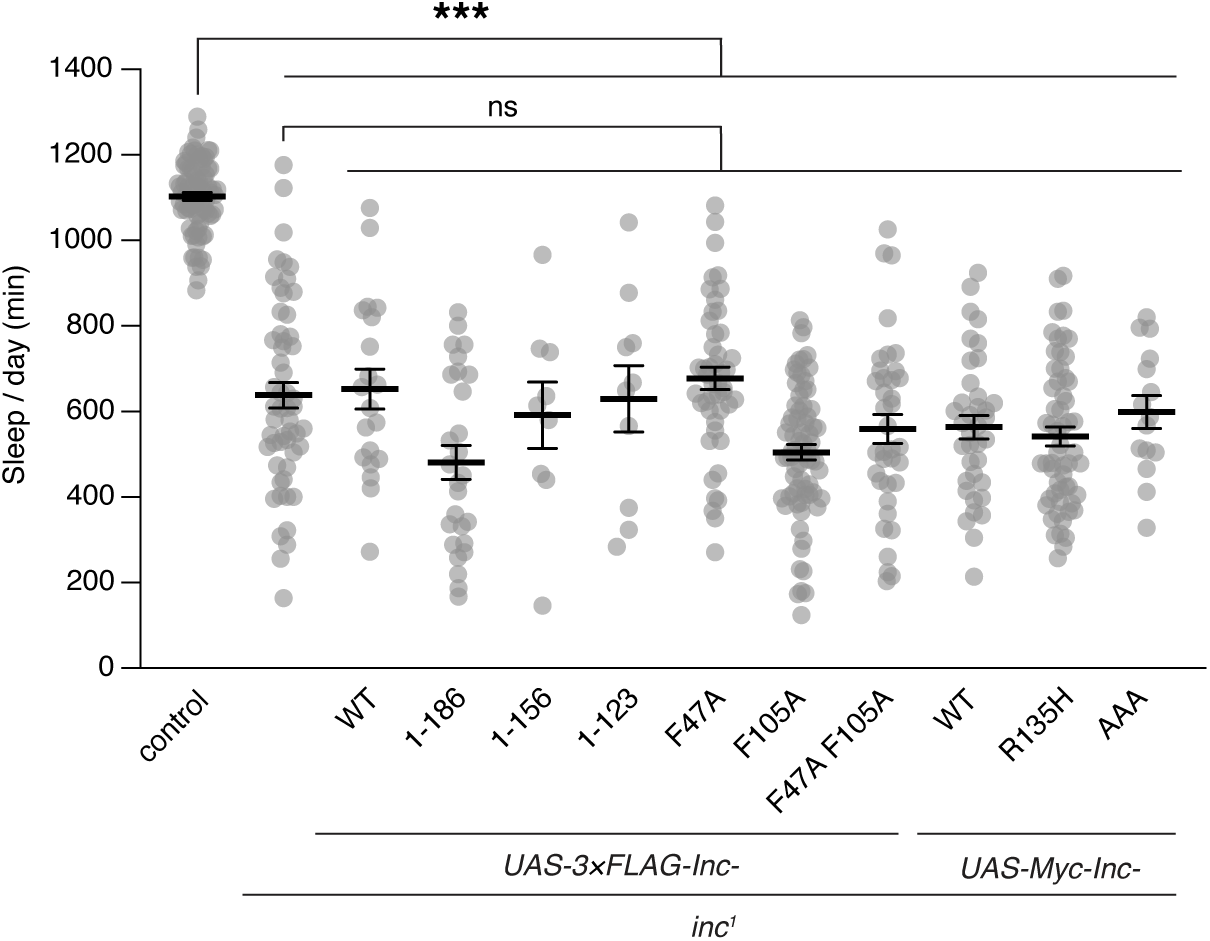
*UAS-Inc* transgenes do not alter sleep in *inc* mutants in the absence of Gal4. Total daily sleep is shown for control, *inc^1^*, and *inc^1^* animals heterozygous for indicated UAS transgenes. Mean ± SEM is shown. n = 83-65; Kruskal-Wallis and Dunn’s tests; ***p < 0.001; ns, not significant (p > 0.05).

**S9 Fig.**
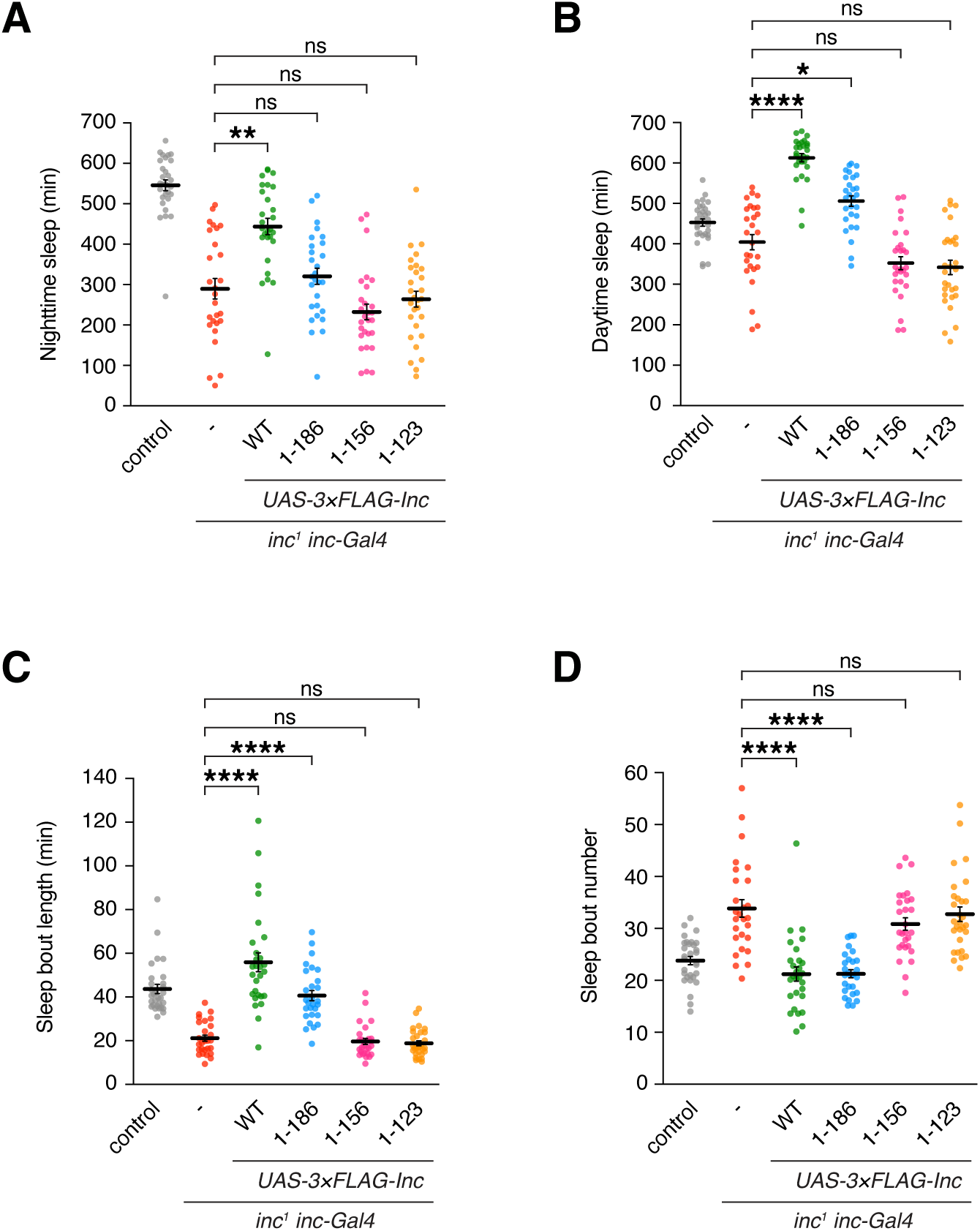
Additional sleep parameters for animals expressing Inc C-terminal truncations. **(A-D)** Sleep parameters for *inc^1^ inc-Gal4* animals expressing 3×FLAG-tagged Inc or C-terminally truncated Inc mutants. **(A)** Nighttime sleep. **(B)** Daytime sleep. **(C)** Sleep bout duration. **(D)** Sleep bout number. Mean ± SEM is shown. n = 27-30 as in Fig 4B; Kruskal-Wallis and Dunn’s tests; *p < 0.05; **p < 0.01; ****p < 0.0001; ns, not significant (p > 0.05). For all panels, animals are heterozygous for UAS transgenes.

**S10 Fig.**
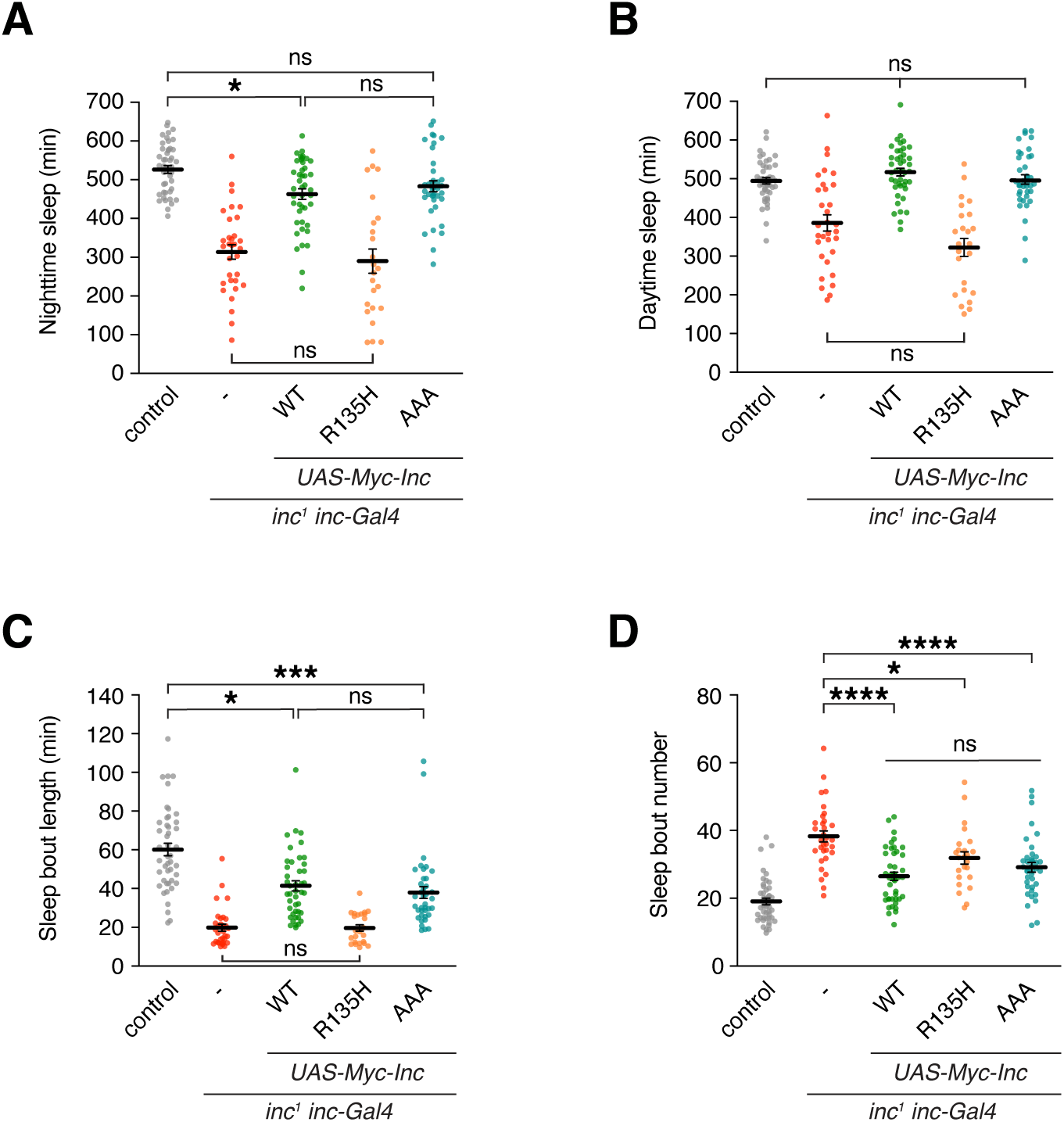
Additional sleep parameters for animals expressing Inc C-terminal point mutants. **(A-D)** Sleep parameters for *inc^1^ inc-Gal4* animals expressing 3×Myc-tagged Inc or Inc point mutants. **(A)** Nighttime sleep. **(B)** Daytime sleep. **(C)** Sleep bout duration. **(D)** Sleep bout number. Mean ± SEM is shown. n = 24-45 as in Fig 6E; Kruskal-Wallis, p<0.0001 and Dunn’s tests; *p < 0.05; ***p < 0.001; ****p < 0.0001; ns, not significant (p > 0.05). For all panels, animals are heterozygous for UAS transgenes.

**S1 Table.** Summary of Inc point mutants targeting Inc-Cul3 interactions.

| **Inc mutant** | **Inc-Inc binding** | **Inc-Cul3 binding** | **Inc stability** |
| --- | --- | --- | --- |
| F47A | nc | strongly reduced | nc |
| R50E | reduced | reduced | nc |
| D57A | nc | nc | nc |
| D61A | nc | nc | nc |
| E104K | nc | nc | nc |
| F105A | nc | strongly reduced | nc |
| Y106F | nc | nc | nc |
| N107A | nc | nc | nc |
| F47A F105A | nc | strongly reduced | nc |

nc, no change

**S2 Table.** Summary of Inc point mutants targeting Inc-Inc interactions. (PDF)

| **Inc mutant** | **Inc-Inc binding** | **Inc-Cul3 binding** | **Inc stability** |
| --- | --- | --- | --- |
| T36A | nc | nc | nc |
| T36A D71A | nc | nc | nc |
| D71A R85E | N/A | N/A | strongly reduced |
| D73A N82A | nc | nc | nc |
| K88D E101K | nc | reduced | nc |
| T36A D71A R85E | N/A | N/A | strongly reduced |

nc, no change
N/A, not able to be assessed due to instability

**S1 Data. Source data for figures and supplemental figures.**

**(XLSX)**

